# Lipid Chaperoning of a Thylakoid Protease Whose Stability is Modified by the Protonmotive Force

**DOI:** 10.1101/788471

**Authors:** Lucas J. McKinnon, Jeremy Fukushima, Kentaro Inoue, Steven M. Theg

## Abstract

Protein folding is a complex cellular process often assisted by chaperones but can also be facilitated by interactions with lipids. Disulfide bond formation is a common mechanism to stabilize a protein. This can help maintain functionality amidst changes in the biochemical milieu which are especially common across energy-transducing membranes. Plastidic Type I Signal Peptidase 1 (Plsp1) is an integral thylakoid membrane signal peptidase which requires an intramolecular disulfide bond for *in vitro* activity. We have investigated the interplay between disulfide bond formation, lipids, and pH in the folding and activity of Plsp1. By combining biochemical approaches with a genetic complementation assay, we provide evidence that interactions with lipids in the thylakoid membrane have chaperoning activity towards Plsp1. Further, the disulfide bridge appears to prevent an inhibitory conformational change resulting from proton motive force-mimicking pH conditions. Broader implications related to the folding of proteins in energy-transducing membranes are discussed.

## Introduction

Biological membranes form the boundary between cells and their external environment and are responsible for compartmentalization in eukaryotic cells. Known to be much more than a passive barrier, membranes carry out a wide range of essential processes for living cells, including generation and maintenance of electrochemical gradients, perception and transduction of internal and external signals, transport of biomolecules, and synthesis of lipids. Each of these diverse biological processes requires integral and peripherally associated membrane proteins. Due to their tight association with membranes, the structures and functions of membrane proteins can be influenced by properties of the bilayer such lipid composition, fluidity, charge, and thickness. Indeed, there is a vast amount of evidence supporting a crucial role of the lipid bilayer in the function and/or regulation of membrane proteins. First, phospholipids are present in the structures of numerous membrane proteins and complexes (Lee, 2011) including photosystem II (PSII) from cyanobacteria (Umena et al., 2011) and spinach (Wei et al., 2016). In many cases, lipids are well-resolved in protein crystal structures and bind in specific grooves, analogous to prosthetic groups (Lee, 2004). Second, specific lipid species are required for the activity (Lee, 2004; Soom et al., 2001), proper folding (Dowhan et al., 2004), or membrane insertion (van Klompenburg et al., 1998) of a variety of membrane proteins. Finally, membrane lipids can influence transmembrane helix packing (Lee, 2011), induce conformational changes of extra-membrane domains (Hansen et al., 2011), or stimulate oligomerization of membrane proteins (Stangl and Schneider, 2015). These examples highlight the remarkable versatility of membrane lipids.

In addition to lipids, a fundamental part of the molecular environment of proteins within energy-transducing membranes is the proton motive force (pmf) consisting of both a proton gradient (ΔpH) and an electrical gradient (Δψ). Changes in the magnitude of the pmf and/or the partitioning between Δψ and ΔpH have the potential to alter protein structure and function in several ways, for instance, alteration of the protonation state and in turn the polarity of ionizable side chains (Hamsanathan and Musser, 2018). Consistent with this idea is the observation of pmf-driven conformational changes in voltage-gated ion channels (Catterall et al., 2017) as well as pmf-dependent oligomerization of TatA, a membrane protein involved in the transport of folded proteins in bacteria and chloroplasts via the twin-arginine translocon (Hamsanathan and Musser, 2018). Thus, the structure and function of membrane proteins can be influenced not only by lipids in a bilayer but also by a pmf.

Thylakoid membranes are the energy-transducing membranes in chloroplasts and cyanobacteria and are replete with proteins whose structure and function are influenced by lipids and/or a pmf. For example, the structural stability of light-harvesting LHCII trimers in proteoliposomes is significantly enhanced by the galactolipid monogalactosyl diacyl glycerol (MGDG) (Seiwert et al., 2017), which comprises up to 50% of the total lipids found in thylakoid membranes (Dormann and Benning, 2002). Another example is the membrane protein PsbS which is influenced by both lipids and the pmf. Specifically, insertion of denatured PsbS into liposomes made of thylakoid lipids promotes it to fold into the native conformation (Liu et al., 2016). Furthermore, protonation of two conserved glutamate residues in PsbS in response to a certain magnitude of ΔpH appears to trigger conformational changes leading to its activation (Niyogi et al., 2005). A third example is the thylakoid lumen protein violaxanthin deepoxidase (VDE), in which the catalytic activity (Jahns et al., 2009) and reversible membrane association (Hager and Holocher, 1993) is ΔpH-dependent. VDE activity is also stimulated in the presence of liposomes containing MGDG (Latowski et al., 2002) and by exogenous MGDG (Latowski et al., 2000). Although not an integral membrane protein, VDE still responds to the pmf across thylakoids and depends on specific lipids within the membrane. These and other examples make clear the fact that the pmf can serve a broader role than simply providing the energy for ATP synthesis.

The vast majority of chloroplast proteins are encoded in the nuclear genome, synthesized on cytosolic ribosomes, and are post-translationally targeted to the chloroplast (Shi and Theg, 2013). All known soluble thylakoid lumen proteins (Schubert et al., 2002) and several thylakoid membrane proteins (Mant et al., 1994; Michl et al., 1994; Rodrigues et al., 2011) are synthesized as precursors containing bipartite N-terminal transit peptides consisting of a stromal targeting domain and a thylakoid transfer signal (TTS) in tandem. Homologous to bacterial signal peptides (Paetzel et al., 2002), the TTS targets the protein to the thylakoid and is cleaved off on the lumenal side by the integral membrane thylakoidal processing peptidase (TPP), yielding a mature protein (Albiniak et al., 2012). The major isoform of TPP called Plastidic type I signal peptidase 1 (Plsp1) and its homolog LepB1 are essential for photoautotrophic growth and processing of lumen-targeted proteins in plants and cyanobacteria, respectively (Hsu et al., 2011; Shipman-Roston et al., 2010; Zhbanko et al., 2005). Additionally, Plsp1 can cleave the TTS from numerous TPP substrates in Triton X-100 micelles (Midorikawa et al., 2014). Lipids and/or the pmf likely influence the structure and activity of Plsp1 given that (i) Plsp1 exhibits a tendency to bind to thylakoid membranes *in vitro* (Endow et al., 2015), (ii) the majority of the protein resides in the lumen (Midorikawa et al., 2014) in which the pH fluctuates during photosynthesis (Shikanai and Yamamoto, 2017), (iii) it cleaves signal peptides at or near the surface of the membrane (Dalbey et al., 2012), and (iv) exogenous lipids stimulate the activity of the bacterial Plsp1 homolog leader peptidase (LepB) (Tschantz et al., 1995).

Previous biochemical analyses revealed a key role of disulfide bond formation in the structure and function of Plsp1 (Midorikawa et al., 2014). An angiosperm-specific Cys pair (C166 and C286) residing in the lumenal domain of Plsp1 form an allosteric disulfide bond required for Plsp1 *in vitro* activity (Midorikawa et al., 2014). Using a homology-based structural model, disulfide bond reduction was suggested to alter the conformation of Plsp1 leading to the observed loss of activity (Midorikawa et al., 2014). Thus, it was concluded that disulfide bond formation, often referred to as oxidative folding (Kieselbach, 2013), plays a key role in maintaining the active conformation of Plsp1 (Midorikawa et al., 2014). This was interesting given that *E. coli* LepB lacks the corresponding Cys pair (Midorikawa et al., 2014) and contains another non-conserved Cys pair that are dispensable for activity (Sung and Dalbey, 1992).

In this study, we originally sought to confirm that oxidative folding is required for Plsp1 to function *in vivo* and subsequently determine its biological role. Using a genetic complementation assay and a variety of biochemical approaches, we instead discovered that the disulfide bond in Plsp1 is not essential *in vivo* and that membrane lipids alone facilitate the proper folding and, in turn, activity of Plsp1. We also provide evidence that, while the disulfide bridge is not required for Plsp1 activity *in vivo*, it assists in the maintenance of catalytic activity in a relatively acidic environment such as that found in the thylakoid lumen during illumination. Our results present Plsp1 as a novel example of a membrane protein for which lipids act as folding chaperones. In addition, Plsp1 represents the first example, to our knowledge, of a protein that must maintain activity in both stabilizing and destabilizing environments established by the cycle of diurnal energization of the thylakoid membrane. Broader implications related to the folding and homeostasis of proteins in energized membranes are discussed.

## Results

### Disulfide bond reduction causes a conformational change in Plsp1

Prior to this study, we lacked direct evidence that the loss of activity observed *in vitro* after disulfide bond reduction is related to structural changes within Plsp1. Two approaches were taken to investigate this issue.

Our first approach compared the susceptibility of the oxidized and reduced forms of Plsp1 to the protease thermolysin. DTT could not be used as the reductant in this experiment since it inhibited the protease activity of thermolysin (Fig. S1) due to its ability to chelate the critical Zn^2+^ cofactor (Cornell and Crivaro, 1972; Pretzer et al., 1992). Instead, TCEP (tris-2-carboxyethylphosphine), which also diminishes the activity of Plsp1 *in vitro* (Fig. S1), was used as the reductant. Plsp1 extracted from pea thylakoid membranes with Triton X-100 was treated with or without TCEP, followed by incubation with or without thermolysin. Sample aliquots taken over time were examined by SDS-PAGE and immunoblotting using an antibody raised against the pea ortholog of Plsp1 (Midorikawa et al., 2014). The extracted form of Plsp1 remained relatively stable after 30 minutes at room temperature regardless of TCEP treatment (Fig. 1A). In the presence of thermolysin, Plsp1 was degraded in a time-dependent manner, and this degradation was accelerated if Plsp1 was pre-treated with TCEP (Fig. 1A). TCEP-treated Plsp1 migrated more slowly than un-treated Plsp1 on non-reducing SDS-PAGE (Fig. 1B), confirming complete reduction of the disulfide bond (Midorikawa et al., 2014).

**Figure 1:**
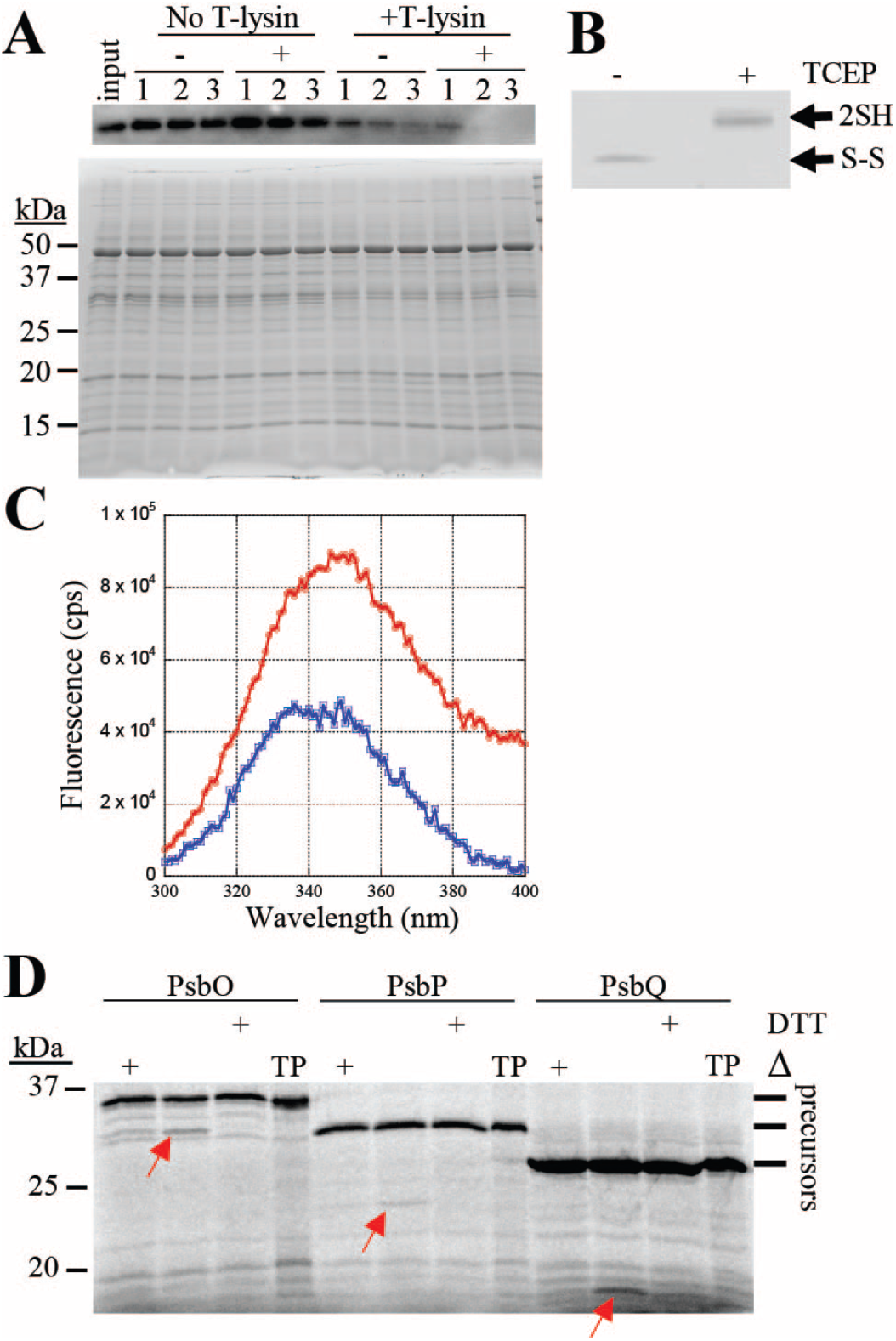
Disulfide bond reduction alters the structure of Plsp1. **(A)** Proteins extracted from Pea thylakoids with 0.25% v/v Triton X-100 were pre-treated with (+) or without (-) 10 mM TCEP followed by incubation with or without thermolysin. Aliquots taken at 10 (1), 20 (Boomer et al.), and 30 (3) minutes were analyzed by SDS-PAGE and immunoblotting with an antibody against Pea Plsp1 (upper panel). A second gel was loaded with half of the amount of each sample and stained with Coomasie Brilliant Blue after electrophoresis (lower panel). Input = un-treated thylakoid extracts. **(B)** An aliquot of the +/− TCEP pre-treated thylakoid extracts was analyzed by non-reducing SDS-PAGE and immunoblotting with the antibody against Pea Plsp1. 2SH = reduced Plsp1. S-S = oxidized Plsp1. **(C)** Emission spectrum of purified T7-Plsp1_71-291_ (~0.1 μM) in 1% w/v octyl glucoside after pre-treatment with (Blue) or without (Red) 50 mM DTT plotted as fluorescence in counts per second (cps) versus wavelength. Shown are the spectra after subtracting those of buffer blanks. The excitation wavelength was 285 nm. **(D)** Processing activity of purified T7-Plsp1 against prPsbO, prPsbP, or prPsbQ. T7-Plsp1 used in panel C was mixed with ^35^S-Met-labeled substrates after being boiled for 10 minutes (Δ) or pre-treated with 50 mM DTT on ice for 20 minutes. Reaction mixtures were incubated at ~25° C for 30 minutes and analyzed by SDS-PAGE and autoradiography. Arrows denote the processed forms of each substrate tested. TP = translation products.

Our second approach to relate Plsp1’s disulfide to its conformation was based on the intrinsic fluorescence of the three native Trp residues in a purified form of Plsp1 containing an N-terminal T7 tag. When excited with 285nm light, Plsp1 exhibited strong fluorescence in the range of 300-400nm with a λ_max_ of 350 ± 2nm (Fig. 1C), which is characteristic of a solvent-exposed Trp side chain. After treatment with DTT, the fluorescence spectrum of Plsp1 exhibited a similar shape and λ_max_ (349 ± 2nm) but with a significant decrease in emission intensities (Fig. 1C). The decrease in fluorescence intensity cannot be attributed to denaturation and aggregation of Plsp1 since the DTT-treated enzyme remains completely soluble (Fig. S1). Since the quantum yield often decreases when a Trp side chain shifts from a buried to a more solvent-exposed local environment (Teale, 1960), reducing the disulfide bond in Plsp1 likely causes a conformational change which increases the exposure of one or more Trp residues to the bulk solvent. As expected, the activity of the enzyme used for the fluorescence experiments is abolished by DTT (Fig. 1D). Taken together, the results presented in Fig. 1 provide direct evidence that the loss of Plsp1 activity upon disulfide bond reduction can be attributed to a conformational change to an inactive structure.

### Plsp1 Cys form a disulfide bond in intact chloroplasts

Previous work showed that Cys166 and Cys286 can form an intramolecular disulfide bond in a recombinant form of Plsp1 as well as in Plsp1 in isolated chloroplasts from pea and Arabidopsis (Midorikawa et al., 2014). To confirm that this disulfide bond forms *in vivo*, we analyzed the *in vivo* redox state of the Cys in Plsp1 using a thiol labeling assay (Shapiguzov et al., 2016). The endogenous Plsp1 protein in Arabidopsis plants could not be monitored due to a low level of expression. Instead, we transiently expressed T7-tagged Plsp1 in *Nicotiana benthamiana* leaves to achieve high expression and facilitate detection. A genetic complementation experiment confirmed that T7-Plsp1 is indeed functional (Fig. S2).

Intact chloroplasts were isolated from leaves after infiltration with Agrobacterium containing a plasmid encoding T7-Plsp1 or after a mock infiltration and were used to probe the redox state of the two Cys residues in Plsp1. After blocking free thiol groups with N-ethyl maleimide (NEM), proteins were treated with TCEP followed by addition of methoxypolyethylene glycol maleimide (mPEG-MAL), which confers a large size shift upon labeling the newly-exposed thiols. A portion of T7-Plsp1 displays a mobility shift from ~27-kDa to ~50-kDa when TCEP treatment precedes labeling with mPEG-MAL (Fig. 2A). This size shift is comparable to that observed when both Cys residues of *in vitro* translated Plsp1 are labeled with mPEG-MAL (Fig. S3). We therefore interpret the 50-kDa band as T7-Plsp1 with mPEG-MAL on both Cys residues. This size shift was detected regardless of NEM treatment (Fig. 2A) which is expected if Plsp1 is initially disulfide bonded. As a control, the single Cys residue in OE23 was labeled by mPEG-MAL but only when NEM was omitted from the initial chloroplast lysis and membrane solubilization (Fig. 2A) confirming that the NEM successfully blocked free thiols in the initial steps of the experiment.

**Figure 2.**
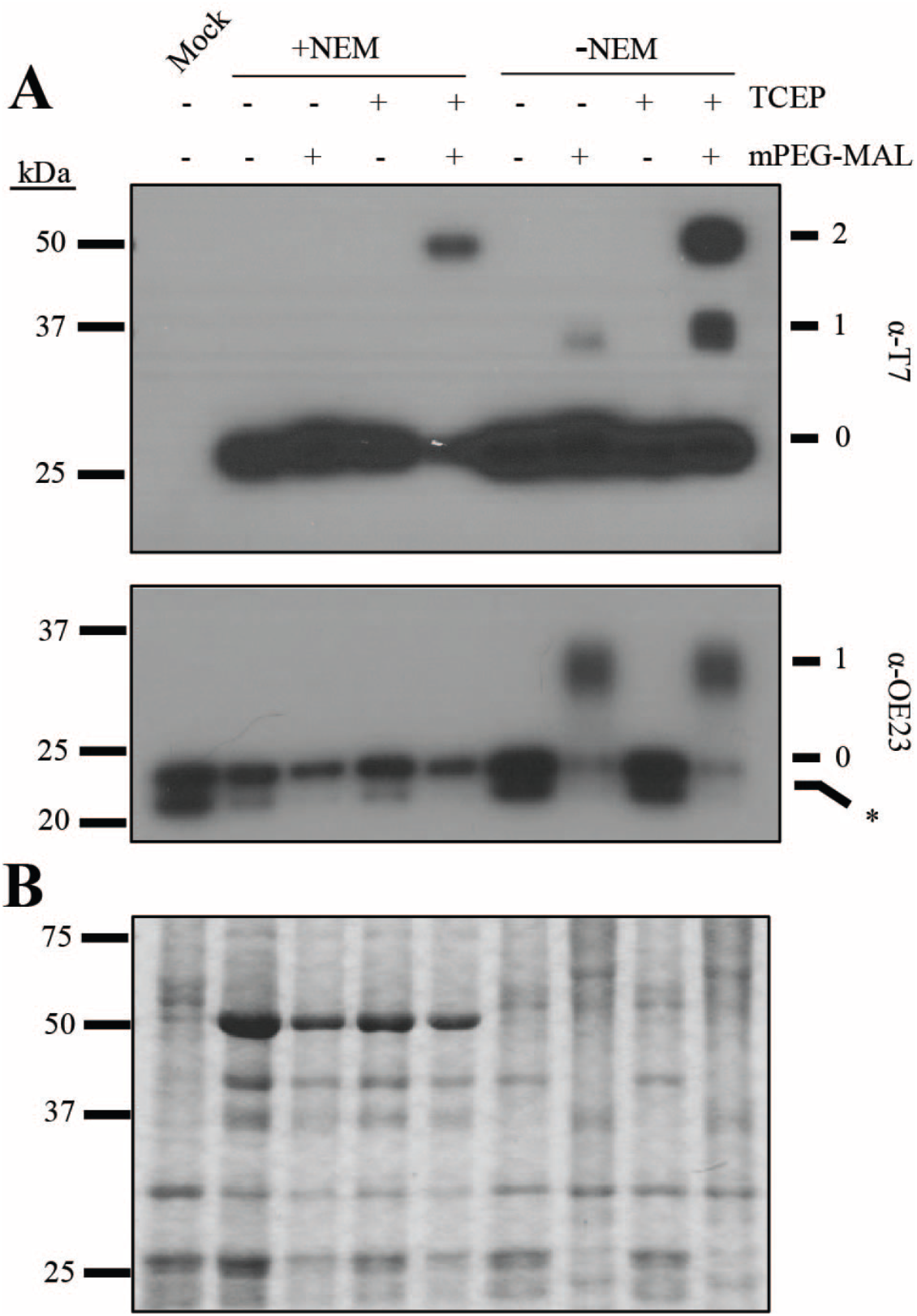
Plsp1 Cys form a disulfide bond in chloroplasts. **(A)** PEG-MAL labeling of Cys in isolated chloroplast membranes. Thiol labeling was performed as described in Materials and Methods. Samples were analyze by SDS-PAGE and immunoblotting using the α-T7 antibody for detection of T7-Plsp1 (top panel) or the α-OE23 antibody as a control (bottom panel). Unlabeled protein (0). 1 or 2 Cys residues labeled with mPEG-MAL, (1) and (Boomer et al.), respectively. Asterisk indicates unknown immunoreactive protein in *N. benthamiana* chloroplast membranes. **(B)** Coomasie-stained gel of the same samples analyzed in figure A

A minor proportion of T7-Plsp1 was shifted to ~37-kDa (Fig. 2A) after TCEP + mPEG-MAL treatment. In addition, a significant proportion of T7-Plsp1 was unlabeled even without NEM treatment. We attribute these results to incomplete mPEG-MAL labeling of thiols and/or disulfide bond reduction by TCEP, and we interpret the ~37-kDa band as T7-Plsp1 with mPEG-MAL on one Cys because the mobility is similar to that of the labeled forms of *in vitro* translated single Cys Plsp1 mutants (Fig. S3), Regardless of the technical limitations, it is clear that a significant portion of the T7-Plsp1 population in intact chloroplasts contains an intramolecular disulfide bond.

### Processing activity is required for the in vivo function of Plsp1

Lack of Plsp1 leads to defective thylakoid development as well as the accumulation of unprocessed forms of several thylakoid lumen proteins (Midorikawa and Inoue, 2013; Shipman-Roston et al., 2010). In addition, overexpression of either of the other two Plsp isoforms (i.e., Plsp2A and Plsp2B) does not rescue the *plsp1-1* T-DNA knockout mutant (Hsu et al., 2011).

These findings led to the conclusions that (i) Plsp1 is the major isoform of the thylakoidal processing peptidase and (ii) Plsp1 is responsible for cleaving most, if not all, TTSs in chloroplasts. To confirm that Plsp1 directly cleaves thylakoid-transfer signals *in vivo*, we tested whether changing the catalytic nucleophile Ser142 (Midorikawa et al., 2014) to Ala (i.e., S142A) would disrupt the function of Plsp1 using a genetic complementation approach, as described previously (Endow and Inoue, 2013). For two independent S142A lines, we observed albino hygromycin-resistant seedlings exhibiting a stunted growth phenotype similar to the *plsp1-1* mutant (Fig. S4). Genomic PCR confirmed that the albino hygromycin-resistant seedlings carried the *CITRINE-PLSP1* transgene in the *plsp1-1* mutant background (Fig. S4). SDS-PAGE and immunoblotting analysis of total seedling extracts revealed that the Citrine-Plsp1 protein was synthesized in each of the transgenic lines (Fig. S4). Furthermore, PsbO and Toc75 both accumulated as unprocessed forms in the albino S142A seedlings as in the *plsp1-1* mutant (Fig. S4). Taken together, these data show that expression of a non-catalytic Plsp1 variant cannot rescue the *plsp1-1* mutant providing a necessary control for the experiments described in Fig. 3.

**Figure 3.**
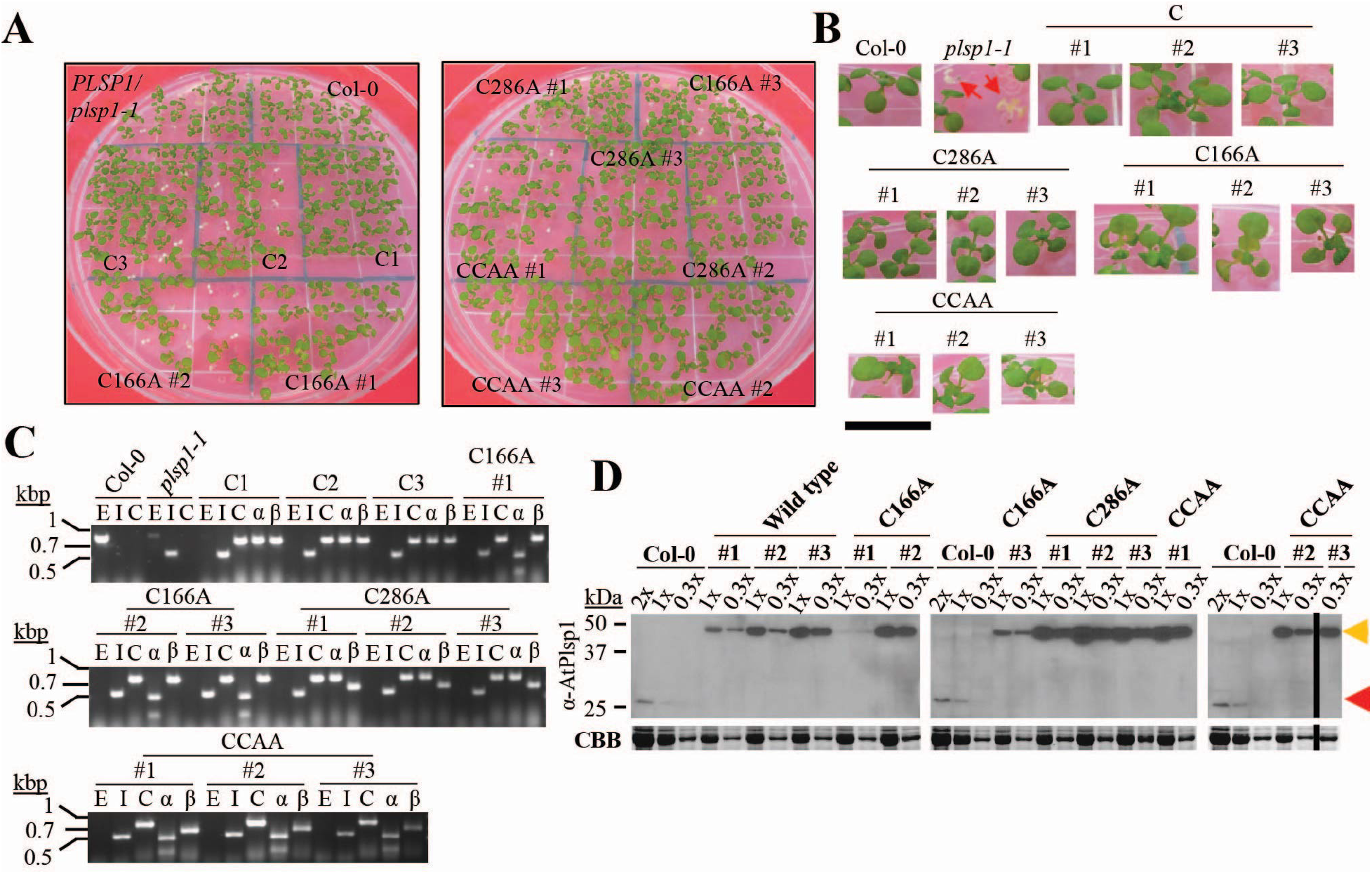
Expression of redox-inactive Citrine-Plsp1 variants rescues the *plsp1-1* knockout mutant. **(A)** Images of 12 day-old seedlings growing on MS medium supplemented with 1% w/v sucrose. *C = CITRINE-PLSP1, C166A* = *CITRINE-PLSP1-C166A*, C286A = *CITRINE-PLSP1-C286A*, CCAA = *CITRINE-PLSP1-C166A/C286A*. **(B)** Close-up images of seedlings from panel A. Red arrows indicate *plsp1-1* null mutants. Scale bar represents 1cm. **(C)** Results of genomic PCR experiment to confirm seedling genotypes. E = *PLSP1*, I = *plsp1-1*, C = *CITRINE-PLSP1*, α = *PLSP1-C166A* (PstI digestion of C), β = *PLSP1-C286A* (PvuII digestion of C). **(D)** Expression of Citrine-Plsp1 proteins in isolated chloroplasts. Proteins from isolated chloroplasts were analyzed by SDS-PAGE and immunoblotting using the α-AtPlsp1antibody. RbcL bands on duplicate gels stained with Coomasie Brilliant Blue (CBB) are shown below as a loading control. The vertical black bar separates non-adjacent lanes from the same membrane. 1X =1 μg chlorophyll equivalent. Red and orange arrows indicate Plsp1 and Citrine-Plsp1, respectively.

### Expression of redox-inactive Plsp1 variants rescues the plsp1-1 mutant

To assess whether the *in vivo* activity of Plsp1 requires a disulfide bond, we tested the functionality of Citrine-tagged Plsp1 variants in which one or both Cys are replaced by Ala. Surprisingly, plants carrying a *CITRINE-PLSP1-CA* (*C166A, C286A*, or *C166A/C286A*) transgene in the *plsp1-1* mutant background exhibited a wild type-like phenotype (Fig. 3A and 3B). We isolated three independent complemented lines for each construct and checked for the presence of Cys/Ala mutations in *PLSP1* by a restriction digest of genomic PCR products (Fig. 3C). Chloroplasts from each of the complemented lines expressed Citrine-Plsp1 and lacked the endogenous form of Plsp1 (Fig. 3D). Notably, the expression of the Citrine-Plsp1 protein was highly variable across the transgenic lines ranging from close to endogenous Plsp1 levels (e.g. C166A #1) to a level well above the dynamic range of detection (e.g. C286A #2). In addition, the wild-type Citrine-Plsp1 and endogenous Plsp1 displayed redox-dependent mobility on SDS-PAGE whereas the mobility of the Citrine-Plsp1-CA proteins was not redox-dependent (Fig. S5). This indicates that changing one or both Cys in Plsp1 to Ala prevents disulfide bond formation.

All twelve complemented lines we isolated were homozygous for the *plsp1-1* null allele, but five of them were hemizygous for the *CITRINE-PLSP1* transgene (Fig. S5). For each of these lines, the ratio of hygromycin-resistant to-susceptible seedlings was ~3:1 (Fig. S5), consistent with segregation for a single transgene insertion. Albino hygromycin-resistant seedlings were never observed. Genomic PCR analysis allowed us to determine that the green wild type-like seedlings were complemented, and the albino seedlings were non-transgenic *plsp1-1* mutants. Results of our genetic complementation assays clearly showed that while a disulfide bond is required for *in vitro* activity, it is not essential for Plsp1 activity *in vivo*.

### Wild type and redox-inactive Citrine-Plsp1 variants have comparable activity in chloroplasts

To qualitatively examine the *in vivo* activity of Citrine-Plsp1-CA proteins, we checked the sizes of two Plsp1 substrates in isolated chloroplasts. Chloroplasts from wild-type plants and those from all plants complemented with Citrine-Plsp1 accumulated PsbO and Toc75 in their mature/processed forms (Fig. 4A). Unprocessed forms of these proteins were only detected in total protein extracts from *plsp1-1* mutant seedlings (Fig. 4A).

**Figure 4.**
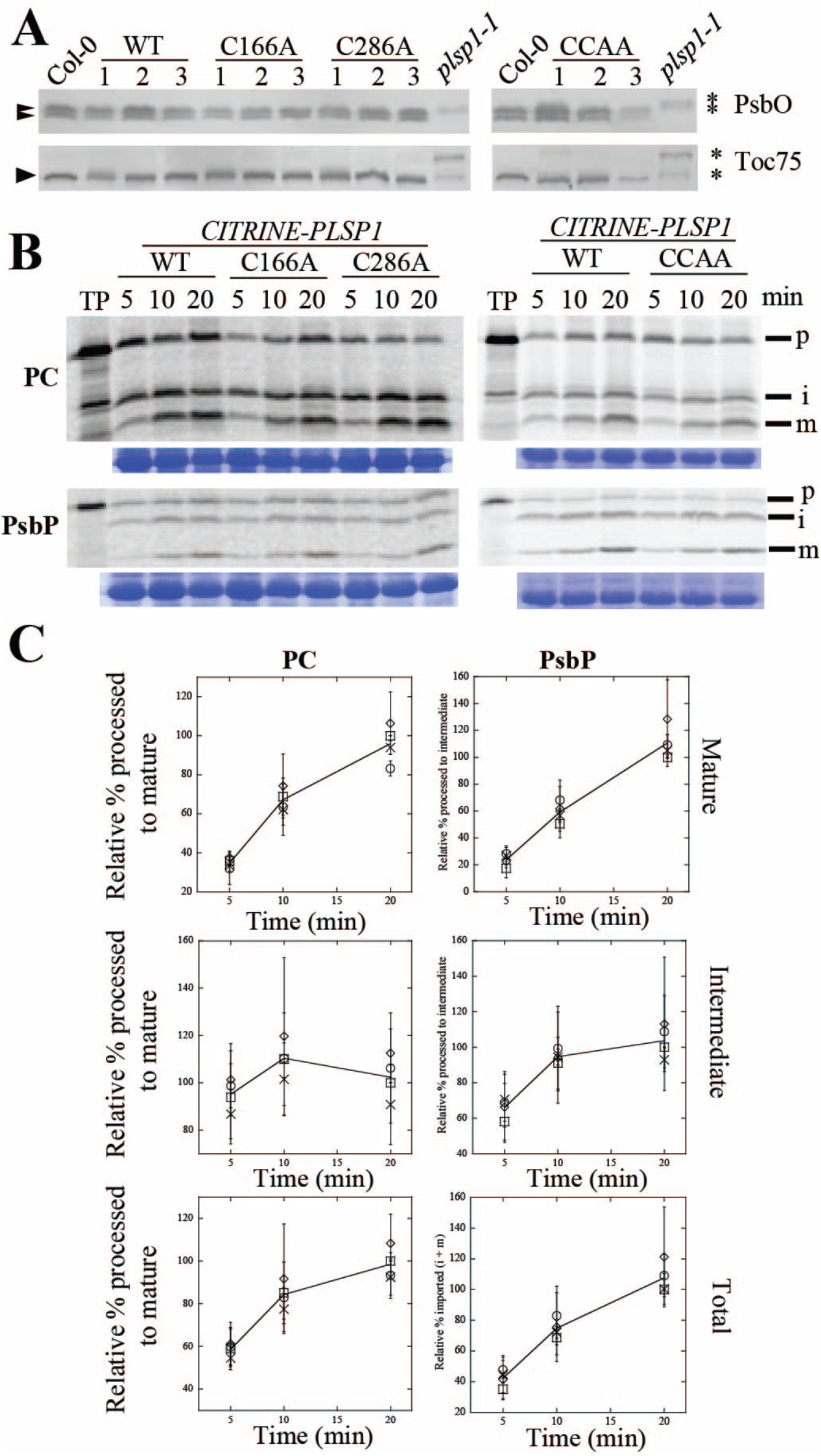
Citrine-Plsp1-CA is active in chloroplasts. **(A)** Size of Plsp1 substrates in chloroplasts isolated from each of the complemented lines shown in Figure 3. Total protein extracts from *plsp1-1* null mutants were also used as a control. Proteins were separated on 12% (α-PsbO) or 7.5% (α-Toc75) SDS-PAGE and detected by immunoblotting using antibodies stated at the right. Asterisks indicate unprocessed forms, and arrowheads indicate processed/mature form (s) of each protein. **(B)** Time course of in vitro import into isolated chloroplasts. ^35^S-Met-labeled forms of each precursor protein were incubated with isolated chloroplasts for 5, 10, or 20 minutes. Intact chloroplasts were re-isolated through a 35% Percoll cushion and washed once in import buffer. A portion of the recovered chloroplasts was used for chlorophyll quantification to normalize gel loading, and the remainder was analyzed by SDS-PAGE and autoradiography. The lines used for the experiments are as follows: C #1, C166A #1, C286A #2, CCAA #2. Each import experiment was repeated three times using chloroplasts isolated on different days. RbcL bands on the CBB-stained gel are shown below as a loading control. P = precursor, i = intermediate, m = mature. PC = plastocyanin. **(C)** Quantification of the import products shown in B. Bands were quantified relative the corresponding wild type band at the 20 minute time point (i.e. wild type maximum). Show are the means ± standard deviation of three biological replicates. The line in each graph is drawn through the average among all four lines at each time point. Squares, wild type; circles, C166A; diamonds, C286A; X, C166A/C286A.

Because most of our complemented lines expressed Citrine-Plsp1 at high levels (Fig. 3D), and high concentrations of Plsp1 exhibit detectable activity despite DTT treatment (Fig. S6), we sought to rule out the hypothesis that the *in vivo* processing activity in chloroplasts containing Citrine-Plsp1-CA mutant proteins is low but sufficient for normal thylakoid development. To that end, we quantitatively examined the activities of each Citrine-Plsp1-CA protein in intact chloroplasts using an *in vitro* protein import assay with radiolabeled precursor proteins. We used the precursors of Arabidopsis PsbP1 and *Silene pratensis* plastocyanin (Last and Gray, 1989) as model substrates for the two thylakoid lumen protein transport pathways cpTAT and cpSEC1, respectively (Albiniak et al., 2012). For time course experiments, we chose one Citrine-Plsp1 wild-type line and one transgenic line representing each Citrine-Plsp1 Cys/Ala mutation. SDS-PAGE and autoradiography of import products showed that all lines imported prPsbP and prPC and processed each to their mature forms at similar rates (Fig. 4B).

Quantification of signals in the autoradiograms revealed that the rate at which mature PsbP or PC accumulate is comparable between each of the CA mutant lines and the wild-type control (Fig. 4C). The total amount of imported PsbP (intermediate + mature) was also comparable over time (Fig. 4C), indicating that chloroplasts from all of the tested lines have similar import efficiencies. Considering that the Citrine-Plsp1 expression level varied between the lines used in this experiment, it appears that the *in vivo* processing rate is not tightly-coupled to the level of Plsp1 in the thylakoid. Based on these data, we conclude that even without a disulfide bond, Plsp1 has a normal level of activity *in vivo*.

### Thylakoid membrane-bound Plsp1 activity is insensitive to DTT

Plsp1 localizes to the thylakoid membrane and presents its catalytic site in the lumen (Kirwin et al., 1988; Shipman and Inoue, 2009). In order to explain its activity *in vivo*, we reasoned that either one or more protein-protein interactions in the thylakoid membrane or interactions with the lipid bilayer compensate for a lack of disulfide bond formation. We developed an assay to directly measure Plsp1 activity in native thylakoids using a membrane permeablization technique (Ettinger and Theg, 1991). Specifically, treatment of isolated thylakoid vesicles with a low concentration of Triton X-100 leads to the release of soluble proteins from the lumen (Ettinger and Theg, 1991). Similarly, we expected that the same low detergent concentration would allow exogenously added substrates to diffuse inside and be processed by Plsp1. Figure 5A (top panel) shows the typical fractionation pattern of proteins from pea chloroplasts after hypotonic lysis followed by treatment of membranes with either a low (0.04%) or high (0.25% v/v) concentration of Triton X-100. The low Triton X-100 treatment released a small amount of the lumenal proteins PsbO and plastocyanin, but Plsp1 remained tightly bound to the membrane (Fig. 5A). In contrast, the high Triton X-100 treatment completely solubilized Plsp1 (Fig. 5A).

**Figure 5.**
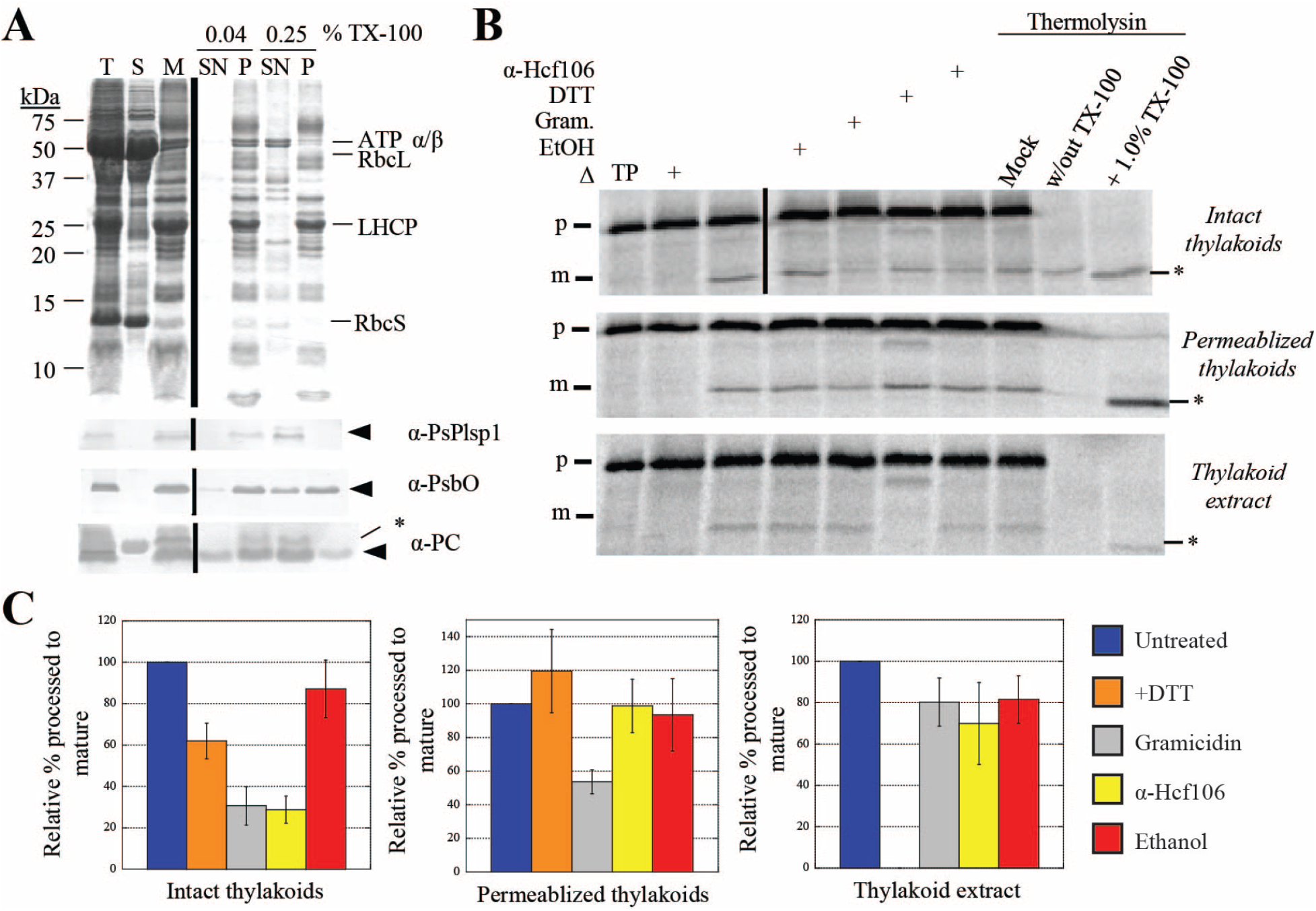
Thylakoid membrane-bound processing activity is insensitive to DTT. **(A)** SDS-PAGE analysis of Pea chloroplast fractions. Proteins were detected by CBB staining (top panel) or immunoblotting (lower three panels) with the antibodies indicated at the right of each panel. Arrowheads indicate proteins of interest, and asterisks indicate non-specific immunoreactive bands. A volume equivalent to 4 μg chlorophyll was loaded in each lane. T = total chloroplasts, S = soluble fraction, M = membranes, SN = supernatant, P = pellet. **(B)** Processing activity in Pea chloroplast membrane fractions depicted in A. 4 μg chlorophyll equivalents of each of the three fractions was pre-treated on ice for 30 minutes with reaction buffer (control), 1% ethanol (EtOH), 30 μM gramicidin (Li et al.) in 1% ethanol, 50 mM DTT, or the α-Hcf106 antibody. As a control, one sample of each chloroplast fraction was heat-inactivated (Δ) at 82°C for 10 minutes prior to adding the substrate. After adding ^35^S-Met-prPsbP, each reaction mixture was incubated in the dark at 28°C for 30 minutes. After the incubation at 28°C, three additional control reactions were treated with 0.5 mM CaCl_2_, thermolysin (0.1 mg/mL with 0.5 mM CaCl_2_), or thermolysin (0.1 mg/mL, with CaCl_2_) in the presence of Triton X-100 (1% v/v) for 40 minutes on ice. Reaction mixtures were quenched by adding an equal volume of 2X sample buffer containing 30 mM EDTA and boiling for 5 minutes. Samples were analyzed by SDS-PAGE and autoradiography. Asterisks indicate a degradation product that was observed in some experiments. **(C)** ImageJ quantification of mature PsbP bands shown in B. Values were normalized to the band intensity in the untreated sample and are expressed as the mean ± standard deviation of at least three independent experiments.

Having established reproducible conditions for preparing membrane-bound and detergent-solubilized forms of Plsp1, we tested the effect of DTT on processing activity in each case using prPsbP as the substrate. Figure 5B shows the results of *in vitro* processing experiments using intact or permeablized (0.04% Triton X-100) thylakoids and thylakoid extract (0.25% Triton X-100 supernatant). The effects of the different treatments on processing activity in intact thylakoids are consistent with proteins that traverse the thylakoid membrane via the cpTAT pathway, of which PsbP is a well-studied substrate (Braun and Theg, 2008). Specifically, both gramicidin and the antibody against Hcf106 (Fig. 5B and 5C) inhibited activity by ~70% due to their effects on protein transport. Gramicidin forms pores in membranes (Kelkar and Chattopadhyay, 2007) which dissipates the pmf on which cpTAT transport depends, and the α-Hcf106 antibody inhibits cpTAT transport by binding to the Hcf106 subunits of the cpTAT translocon (Rodrigues et al., 2011). Interestingly, DTT inhibited processing activity in intact thylakoids by ~40%. It is possible that DTT slightly inhibits the processing activity of Plsp1 in intact thylakoids, but we hypothesize that other non-specific effects are more likely. Resistance to thermolysin confirmed that the processed form of PsbP had been successfully transported into the lumenal compartment of the thylakoid vesicles (Fig. 5B).

Processing activity in permeablized thylakoids relied on simple diffusion of the substrate into the thylakoid vesicles as evidenced by thermolysin susceptibility of processed PsbP and a negligible effect of the α-Hcf106 antibody (Fig. 5B). Importantly, DTT did not affect activity in permeablized thylakoids (Fig. 5B), providing further support that the thylakoid membrane-bound form of Plsp1 is active without a disulfide bond. Gramicidin inhibited processing activity by ~45% (Fig. 5B and 5C) which may be due to unspecified direct or indirect effects on the structure of Plsp1. As expected, the activity in thylakoid extracts was completely inhibited by DTT (Fig. 5B and 5C).

We also used this assay system with Citrine-Plsp1-complemented plants (wild type and C166A/C286A). Permeablized thylakoids from both genotypes processed prPsbP to the mature form, were unaffected by DTT, and were completely inhibited by the type I signal peptidase inhibitor Arylomycin A2 (Paetzel, 2014) (Fig. S7). As expected, processing activity in the wild-type thylakoid extracts was completely inhibited by DTT and Arylomycin A2 (Fig. S7). In comparison, thylakoid extracts containing Citrine-Plsp1-C166A/C286A lacked detectable activity even in the absence of DTT (Fig. S7). The above results (Fig. 5 and Fig. S7), as well as our genetic complementation data (Fig. 3 and Fig. S4), provide compelling evidence that Plsp1 requires a disulfide bond for activity only when it is taken out of its native context.

### Compensation for lack of a disulfide bond in Plsp1 is not due to a stable association with PGRL1

The experiments described in Fig. 5 cannot distinguish between effects of protein-protein interactions or lipid interactions. More than 50% of the total Plsp1 pool in thylakoids forms a stable complex with PGRL1 (Endow and Inoue, 2013), a key component of antimycin A-sensitive cyclic electron flow around photosystem I (Strand et al., 2016). All Plsp1 substrates examined accumulate as their mature forms in chloroplasts lacking PGRL1 (Endow and Inoue, 2013), but Plsp1 is presumably oxidized in the mutant chloroplasts. Two-dimensional blue-native/SDS-PAGE followed by immunoblotting showed that a portion of the Citrine-Plsp1 and PGRL1 pools co-migrate despite changing the Cys in Plsp1 to Ala (Fig. 6A), suggesting that the stable association does not require a disulfide bond in Plsp1. We therefore hypothesized that the interaction with PGRL1 might keep the reduced form of Plsp1 catalytically active.

**Figure 6.**
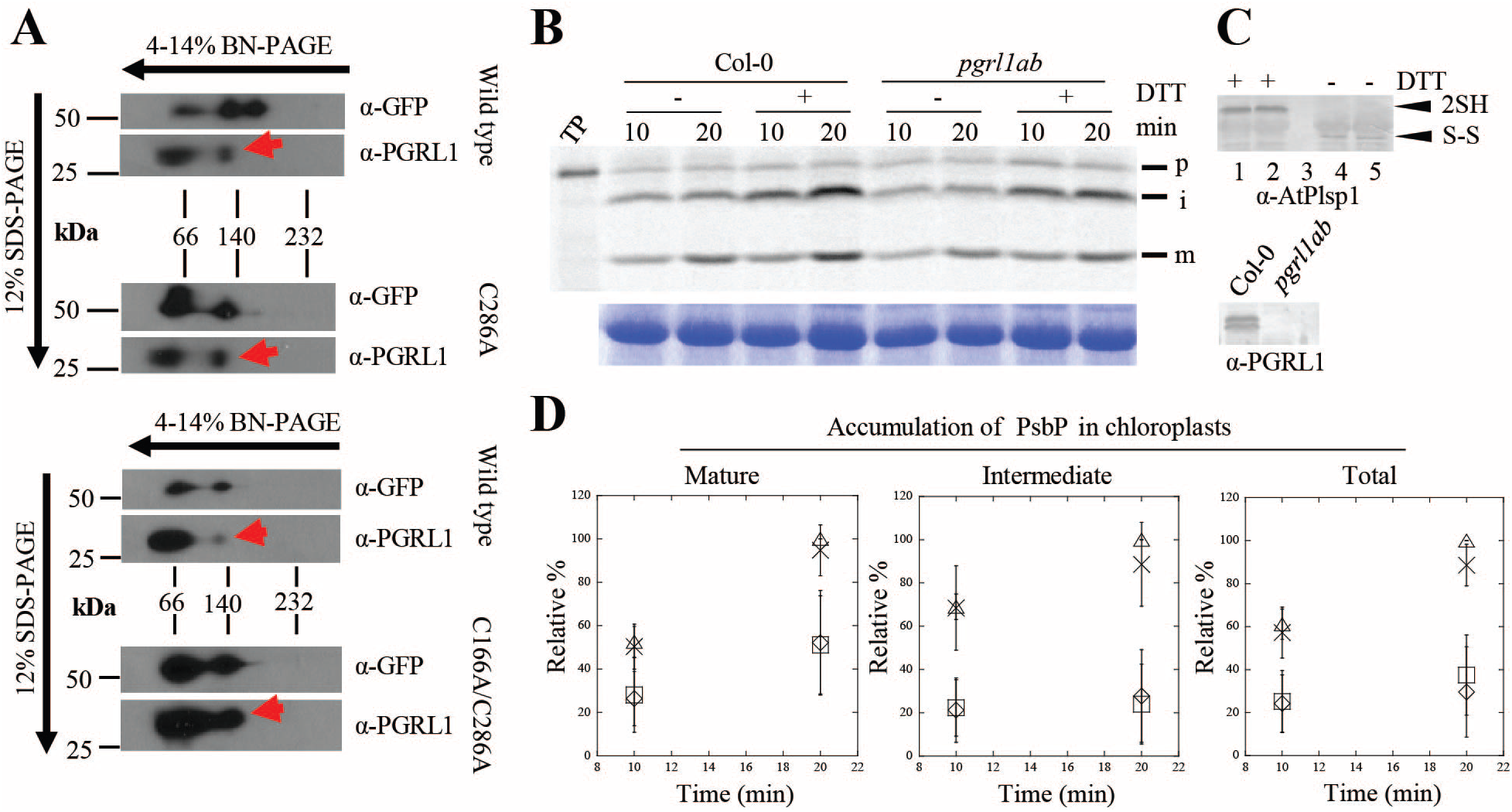
Association with PGRL1 does not compensate for lack of Cys in Plsp1. **(A)** 2D-BN/SDS-PAGE analysis of Arabidopsis chloroplasts complemented with Citrine-Plsp1, Citrine-Plsp1-C286A, or Citrine-Plsp1-C166A/C286A. Total chloroplasts equivalent to 5 μg (left) or 7.5 μg chlorophyll were solubilized with 1.3% w/v n-dodecyl-β-D-maltoside and run on 4-14% BN-PAGE gels. Lanes were then excised, heated at ~80°C for 30 minutes in denaturing buffer (65 mM Tris-HCl pH 6.8, 3.3% SDS, 570 mM β-ME), and overlayed on 12% SDS-PAGE gels. After electrophoresis, proteins were blotted to PVDF membranes. Membranes were cut at ~37-kDa and probed with the GFP antibody (top) or the PGRL1 antibody (bottom). Red arrows indicate the population of PGRL1 that co-migrates with Citrine-Plsp1. **(B)** Analysis of processing activity during in vitro import. Chloroplasts pre-treated on ice for 30 mins with or without 50 mM DTT were mixed with ^35^S-Met-prPsbP and MgATP (3 mM final) and were incubated in the light at 20°C for 10 or 20 minutes. Intact chloroplasts were re-isolated as described in figure 3B and analyzed by autoradiography (top panel). RbcL bands from the CBB-stained gels is shown as a loading control (bottom panel). **(C)** Immunoblotting analysis of chloroplasts used for *in vitro* import experiments. (Upper panel), An aliquot of chloroplasts treated with or without 50 mM DTT prior to *in vitro* import were mixed with non-reducing (no β-ME) sample loading buffer and analyzed by SDS-PAGE and immunoblotting using the antibody against Arabidopsis Plsp1. ”2SH” and ”S-S” indicate the reduced and oxidized forms of Plsp1, respectively. 1 and 4 = Col-0, 2 and 5 = pgrl1ab, 3 = blank. (Lower panel), Chloroplasts used for import experiments were analyzed by SDS-PAGE and immunoblotting using the PGRL1 antibody. **(D)** Quantification of import products shown in C. Bands were quantified relative to the corresponding band (s) in the Col-0 + DTT sample at 20 minutes which was set as 100%. Data are expressed as the mean ± standard deviation of three independent experiments. Squares, wild type - DTT; triangles, wild type + DTT; diamonds, *pgrl1ab* -DTT; X, *pgrl1ab* + DTT.

To test this hypothesis, prPsbP was imported into Col-0 wild type chloroplasts or those from the *pgrl1ab* knock out mutant (Endow and Inoue, 2013) after pre-treatment of chloroplasts with or without DTT. If PGRL1 maintains Plsp1 activity, we expected that PsbP would either (i) accumulate exclusively as the intermediate form or (ii) be processed to the mature form more slowly after treatment with DTT in *pgrl1ab* chloroplasts. However, both the intermediate and mature forms of PsbP accumulated at similar rates in the two genotypes regardless of DTT treatment (Fig. 6B and 6C). The total amount of imported PsbP was also similar between both genotypes (Fig 6C). Immunoblotting confirmed that the *pgrl1ab* mutant chloroplasts lacked detectable PGRL1 protein and that the DTT pre-treatment of chloroplasts completely reduced Plsp1 (Fig. 6D). Notably, DTT pre-treatment increased the total amount of PsbP that was imported (Fig. 6C), an effect first observed for import of the ferredoxin precursor into Pea chloroplasts (Pilon et al., 1992). This is most likely due to reductive enhancement of components of the import apparatus (Bolter et al., 2015). Based on these data, we concluded that the association with PGRL1 does not compensate for the lack of a disulfide bond in Plsp1.

### A Lipid Bilayer Abrogates the Effect of DTT on Plsp1 Activity

Having shown that membrane-embedded Plsp1 is not stabilized against reductants by interaction with PGRL1, we sought to test whether this stability was imparted by the membrane environment itself. As pointed out by Hamsanathan and Musser (2018), it is well known that membrane proteins can lose their activities upon removal from membranes. We asked if this would translate into solubilized Plsp1 maintaining its structure in the presence of reductants upon reconstitution into membranes. As a first attempt to answer this question, we reconstituted purified Plsp1 into lipid vesicles prepared from *E. coli* total lipid extract. As seen in Figure S8, this led to a Plsp1 sample that retained approximately 40% of its activity upon reduction with DTT. Encouraged by these results and recognizing the different chemical natures of the lipids in *E. coli* and thylakoids, we repeated these experiments using liposomes composed of MGDG, DGDG, SQDG, and PG, the four major diacylglycerolipids found in thylakoid membranes (Webb and Green, 1991). After detergent removal, Plsp1 was recovered in the pelleted liposomes and remained so after treatment of the proteoliposomes (PLs) with sodium carbonate (Fig. 7A). Under non-reducing conditions, Plsp1 exhibited a faster migration on SDS-PAGE, indicating the presence of the disulfide bond (Fig. 7A). These results provide evidence that Plsp1 was successfully reconstituted into liposomes and that it was in the expected redox state.

**Figure 7.**
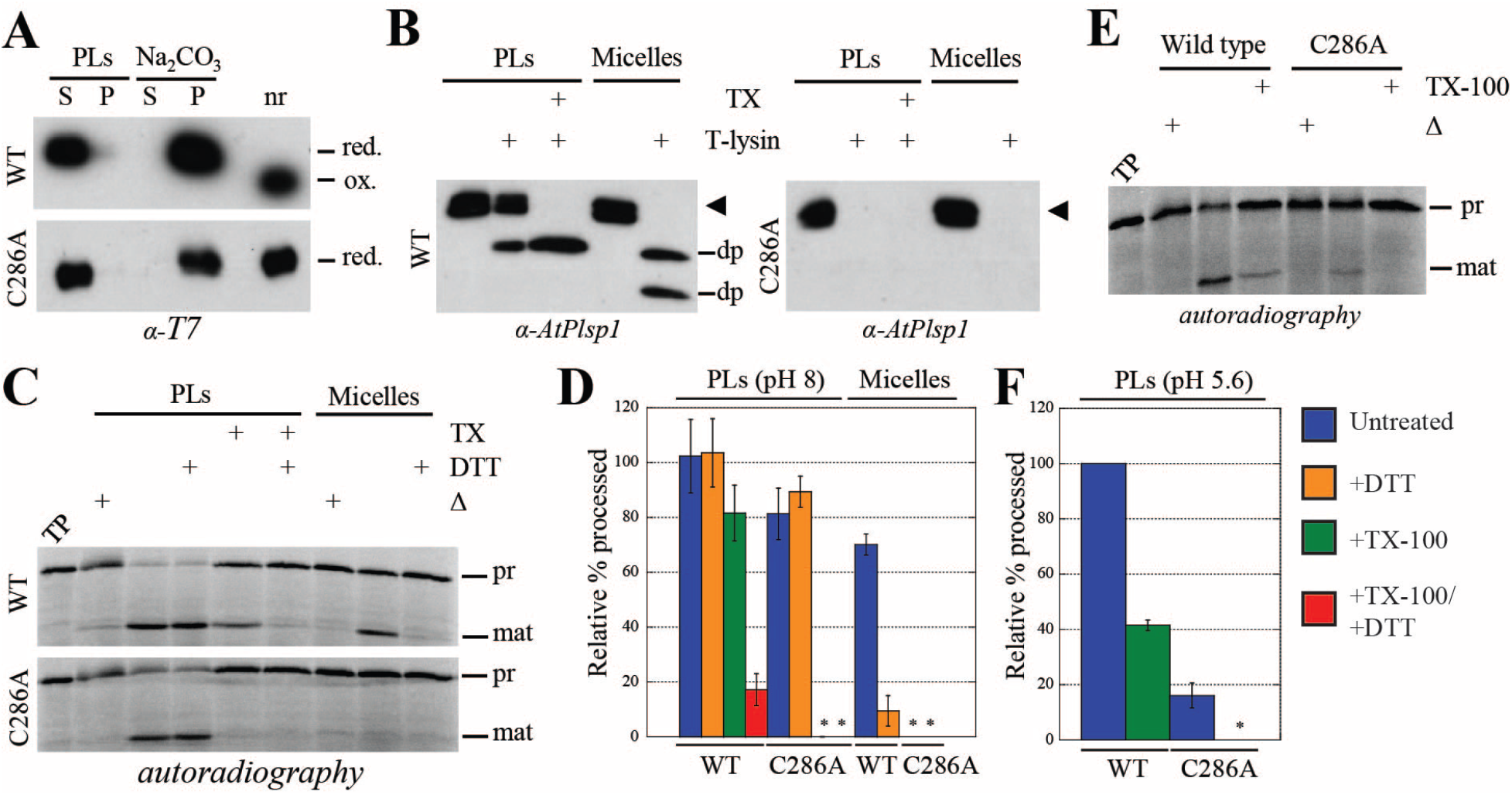
Reconstitution into thylakoid lipid vesicles recovers the activity of redox-inactive Plsp1 in a pH-dependent manner. **(A)** T7-Plsp1 (wild type or C286A) was purified in 0.25% Triton X-100 (micelles, final pH~7.5) and reconstituted into liposomes to yield proteoliposomes (PLs) and were then divided into soluble (S) and pellet (P) fractions by centrifugation as described in Materials in Methods. The soluble fraction was treated with 0.1 M Na_2_CO_3_ for 5 minutes on ice and then subjected to ultracentrifugation to yield soluble and pellet fractions. nr = aliquot run in non-reducing sample buffer. Samples were analyzed by SDS-PAGE and immunoblotting. **(B)** PLs at pH 8 were subjected to treatment with thermolysin (10 μg/mL with 0.5 mM CaCl_2_) with or without 2% Triton X-100 (TX) at 25°C for 40 minutes. Samples were analyzed by SDS-PAGE and immunoblotting. The arrowhead indicates the T7-Plsp1 doublet, and “dp” indicates degradation products observed after thermolysin treatment. **(C)** PLs at pH 8 were pre-treated with or without 50 mM DTT and/or 2% Triton X-100 (TX) for 30 minutes on ice followed by incubation with^35^S-prPsbP for 30 minutes at 25°C. As a control, 60 nM purified T7-Plsp1 pre-treated with or without 50 mM DTT was included. Samples were analyzed by SDS-PAGE and autoradiography. Δ = boiled for 10 minutes prior to adding substrate. TP = translation product. **(D)** Quantification of mature PsbP bands in C relative to the TP on the same gel. Shown are the means ± standard deviation from three independent experiments. WT = wild type. Asterisks indicate undetectable activity. **(E)** PLs resuspended in MES liposome buffer at pH 5.6 were assayed for processing activity. **(F)** Quantification of mature PsbP bands in E relative to wild type -TX-100. Shown are the means ± standard deviation from three independent experiments.

T7-Plsp1 exhibited differential sensitivity to thermolysin in the presence and absence of Triton X-100 (Fig. 7B). In the absence of Triton X-100, a portion of wild-type Plsp1 in PLs was completely resistant to thermolysin, and a portion was truncated to a ~22-kDa degradation product (Fig. 7B, left panel). The degradation product is similar in size to that observed when endogenous Plsp1 is subject to thermolysin treatment (Endow et al., 2015; Shipman-Roston et al., 2010) and likely represents an N-terminal truncation up to the transmembrane region with the catalytic domain of Plsp1 facing inside the vesicle lumen. The portion that is completely resistant to thermolysin likely represents Plsp1 with the tightly-folded oxidized catalytic domain facing the bulk solvent. This is comparable to the observation that Plsp1 can associate with the *cis* face of isolated thylakoids *in vitro* such that the catalytic domain is resistant to thermolysin (Endow et al., 2015). In the presence of Triton X-100, full-length Plsp1 was completely removed by thermolysin, and a similar ~22-kDa degradation product representing the tightly-folded catalytic domain was observed (Fig. 7B, left panel). Purified Plsp1 in 0.25% Triton X-100 was degraded by thermolysin with degradation products of ~22-kDa and ~20-kDa appearing in most experiments (Fig. 7B, left panel). The observation that reconstituted Plsp1 is not completely degraded after bilayer solubilization may be due to differences in Plsp1 folding or thermolysin activity in 2% versus 0.25% Triton X-100, or the presence of lipids in the solubilized proteoliposomes.

We carried out processing assays using reconstituted Plsp1 and purified Plsp1 in Triton X-100 to test whether the lipid bilayer mitigates the effect of DTT on Plsp1 activity. Interestingly, reconstituted Plsp1 exhibited processing activity that was insensitive to DTT (Fig. 7C and 7D). Upon solubilization of PLs with Triton X-100, the processing activity of wild-type Plsp1 was inhibited by DTT to a similar extent as the non-reconstituted form (Fig. 7C and 7D). These results are consistent with those obtained from *in vitro* processing assays using permeabilized thylakoids and thylakoid extracts from the *plsp1-1*-complemented Arabidopsis plants (Fig. S7). These data indicate that integration into liposomes composed of native thylakoid diacyl lipids renders oxidized Plsp1 activity insensitive to reducing agents.

### Chaperoning activity of the lipid environment mediates recovery of activity in non-disulfide bonded Plsp1

We were curious whether the lipid environment only allowed for the maintenance of Plsp1 activity in the presence of reductant, or whether it would mediate the folding of the protein from an inactive conformation. To this end, we reconstituted Plsp1 into PLs after its inactivation by reduction of the detergent-solubilized form. Remarkably, Plsp1 regained significant activity after reconstitution, and this was minimally affected by DTT (Figure S9C and S9D). In addition, this form of Plsp1 was sensitive to thermolysin similar to the oxidized form (Figure S9A) and remained in a reduced state after the reconstitution procedure (Figure S9B). Although solubilization of PLs with Triton X-100 slightly decreased the activity of the oxidized form of Plsp1 (Figure 7D), the decrease in activity was significantly greater for the reduced form of Plsp1 (Figure S9D). Interestingly, DTT inhibited activity when the PLs had been solubilized with Triton X-100 despite the fact the Plsp1 was fully reduced at the end of the reconstitution procedure (Figure S9C and S9D). Our interpretation is that a portion of the reconstituted Plsp1 had become oxidized in between resuspension of PLs in buffer and the addition of substrate (i.e., ~3.5 hours). These results indicate that reduced Plsp1, which had adopted an inactive conformation in its reduced state, had been refolded upon incorporation into PLs, suggesting that the lipid environment provides chaperoning activity directed towards this protein.

We also tested the effect of reconstitution into PLs of the C286A mutant of Plsp1 described above. With one of the two cysteines missing this protein cannot form a disulfide bond, and consequently, displayed no processing activity in its solubilized form (Midorikawa et al., 2014) (see also Fig. S9). As with wild-type Plsp1, the C286A mutant protein was recovered in the PL pellet after reconstitution and was not extracted by carbonate washing (Fig. 7A). As expected, its mobility on SDS-PAGE was not altered by treatment with a reducing agent, indicating a lack of a disulfide bond (Fig. 7A). Remarkably, the reconstituted C286A mutant exhibited processing activity in the PLs, indicating that it was folded by its association with the membrane bilayer. This activity was, of course, insensitive to DTT, but was completely abolished by solubilization of the liposomes with Triton X-100 (Fig. 7C and 7D). In aggregate, our experiments with PLs indicate that integration into liposomes composed of native thylakoid diacyl lipids renders Plsp1 activity insensitive to reducing agents and can also facilitate folding from an inactive to an active conformation in the absence of a disulfide bond.

### The disulfide bridge in Plsp1 may impart conformational stability during the diurnal light/dark cycle

Our experiments suggest that the disulfide bridge that stabilizes the active conformation of Plsp1 may be superfluous while it is in a membrane environment. Yet, the placement of the bridging cysteines is invariant in all the angiosperms (Midorikawa et al., 2014), suggesting some evolutionary pressure to form the disulfide in these plants. We recognized that Plsp1’s environment is not static, and that it likely changes during the day/night cycle of membrane energization. Specifically, the thylakoid membrane goes from a non-energized to an energized state with the imposition of the pmf in the light, and the lumen pH changes from approximately 8 in the dark to ~6 or less under illumination (Kieselbach, 2013). In testing the importance of this, we found that the peptidase activity of T7-Plsp1-C286A in PLs at pH 8 was similar to that of wild-type Plsp1 (Fig. 7C and 7D). When assayed at pH 5.6 however, the activity of the C286A variant in PLs was significantly lower than that of the wild-type and was abolished upon solubilization of the liposomes with Triton X-100 (Fig. 7E and 7F). This suggests that the disulfide bond in Plsp1 may not be required at night (in the dark) but provides structural stability to offset the destabilizing effects of, at the least, the pH component of the pmf developed in the light during the day.

## Discussion

### Plsp1 exhibits a conformational change upon reduction of its disulfide bond in detergent micelles

The observed loss of Plsp1 activity upon reduction of the disulfide bond was ascribed to a conformational change in the enzyme (Midorikawa et al., 2014), consistent with well-documented examples of conformational differences between oxidized and reduced forms of proteins (Choi et al., 2001; Gopalan et al., 2006; Nishii et al., 2015; Tanaka and Wada, 1988). Results of our protease susceptibility assay and tryptophan fluorescence analysis support this hypothesis. Plsp1 became more susceptible to the protease thermolysin after treatment with a reducing agent (Fig. 1A), suggesting that additional protease recognition sites became accessible after the disulfide bond was broken. The significant drop in Trp fluorescence upon treatment with DTT also indicated that at least one of the three Trp residues in Plsp1 had become more solvent-exposed. The fluorescence data likely do not reflect a change in the environment of Trp107 as this residue, which is present just before the transmembrane domain, is located far from the catalytic domain. Further inspection of the predicted structure presented by Midorikawa et al (2014) and Endow et al (2015) revealed that, among all three Trp residues, Trp255 is closest to the active site (~6 Å from the general base Lys192) (data not shown). We therefore speculate that the loss of activity is related to a structural change near Trp255.

### Plsp1 forms a non-regulatory disulfide bond in the thylakoid lumen

Approximately 40% of the confirmed soluble lumen proteome, as well as the lumenal domains of several thylakoid membrane proteins, possess redox-active Cys residues which are either known or predicted to form disulfide bonds (Brooks et al., 2013; Hall et al., 2010; Karamoko et al., 2013; Shapiguzov et al., 2016). The results of our *in vivo* thiol-labeling assay indicate that the two conserved Cys residues in Plsp1 form a disulfide bond in the thylakoid lumen (Fig. 2A), adding it to this growing list. The classic view of disulfide bonds is that they enhance the structural stability of proteins (Betz, 1993; Thornton, 1981), but they can also play a role in allosteric regulation, as is the case for most enzymes involved in carbon fixation (Balsera et al., 2014; Schmidt et al., 2006). Midorikawa et al (2014) hypothesized that the disulfide bond in Plsp1 is reversible *in vivo* and acts to regulate activity similar to the proposed paradigm of lumenal redox regulation (Gopalan et al., 2006; Karamoko et al., 2013; Simionato et al., 2015). However, we do not favor this hypothesis because (i) Plsp1 is functional *in vivo* without a disulfide bond (Fig. 3) and (ii) the activity of Plsp1 in PLs is unaffected by DTT (Fig. 7D).

Oxidation of Plsp1 is expected to occur in the thylakoid lumen due to the nature of its thylakoid targeting mechanism. Thylakoid transport of Plsp1 is mediated by the cpSEC1 pathway (Endow et al., 2015), which can only accommodate unfolded polypeptides (Albiniak et al., 2012). The presence of a disulfide bond between C166 and C286 would introduce a large loop in Plsp1 which would likely make the protein incompatible with the cpSecY translocon. Although some oxidized lumen proteins are transported in a folded conformation by the cpTAT pathway (Hall et al., 2010; Schubert et al., 2002) and may be oxidized in the stroma or the lumen, we predict that membrane translocation must precede oxidation for all cpSEC1 substrates that possess a disulfide bridge.

### Lipids have a major effect on Plsp1 structure and function

Our genetic complementation experiments revealed that, in chloroplasts, Plsp1 is functional without a disulfide bond (Fig. 3), which is in stark contrast to the *in vitro* activity assay in detergent micelles (Fig. 1D and (Midorikawa et al., 2014). Failure of the active site S142A Plsp1 variant to complement the null mutant (Fig. S4) confirmed that processing activity is required for *in vivo* functionality of Plsp1. This also suggested that the other two Plsp isoforms (Plsp2A/2B) do not compensate for a non-catalytic form of Plsp1, which is in line with previous data showing that overexpression of Plsp2A or Plsp2B do not rescue the *plsp1-1* null mutant (Hsu et al., 2011). The surprising result that Plsp1 is catalytically active *in vivo* without a disulfide bond (Fig. 3 and 4A) suggested that Plsp1 activity is sensitive to reducing agents *in vitro* due to a lack of one or more factors which are present in thylakoids. We hypothesized that interactions between Plsp1 and another protein and/or lipids in the membrane prevent the conformational change that leads to a loss of activity.

One of the first pieces of evidence supporting this hypothesis was the observation that the activity of Plsp1 in permeabilized thylakoid vesicles is insensitive to reducing agents (Fig. 5C, 5D, and S7). Additional experiments showed that although the stable association between Plsp1 and PGRL1 is not affected by changing one or both Cys in Plsp1 to Ala (Fig. 6A), this interaction does not have a role in maintaining the active conformation of Plsp1 (Fig. 6B and 6D). In contrast, reconstitution into liposomes rendered Plsp1 activity insensitive to DTT. Further, it restored the activity of previously reduced wild type protein (Figure S9), as well as that of the single Cys Plsp1 mutant (Fig. 7C and 7D). The effect of lipids appears to be completely reversible since solubilization of PLs with Triton X-100 abolished the activity of a reconstituted single Cys Plsp1 (Fig. 7C and 7D) as well as the activity of the Cys-less Plsp1 mutant from isolated thylakoids (Fig. S7). Our interpretation of these data is that interactions between Plsp1 and membrane lipids were necessary to prevent the inhibitory conformational change and were also sufficient to shift the enzyme from an inactive into a catalytically active form (Fig. 8). Activity-inducing conformational changes stimulated by association with membrane lipids are also seen in a class of proteins called bacteriocins (Mel and Stroud, 1993; van der Goot et al., 1991) and in the K^+^ channel Kir2.2 (Hansen et al., 2011). More extreme cases are seen in the thylakoid membrane protein PsbS and in certain bacterial outer membrane β-barrel proteins which can be folded from completely denatured forms into their native conformations in the presence of liposomes (Kleinschmidt, 2015; Liu et al., 2016). Taken together, the above-mentioned results lead us to conclude that thylakoid lipids possess folding chaperone activity towards the lumenal catalytic domain of Plsp1 and that the native folding occurs without, and perhaps prior to, disulfide bond formation.

**Figure 8.**
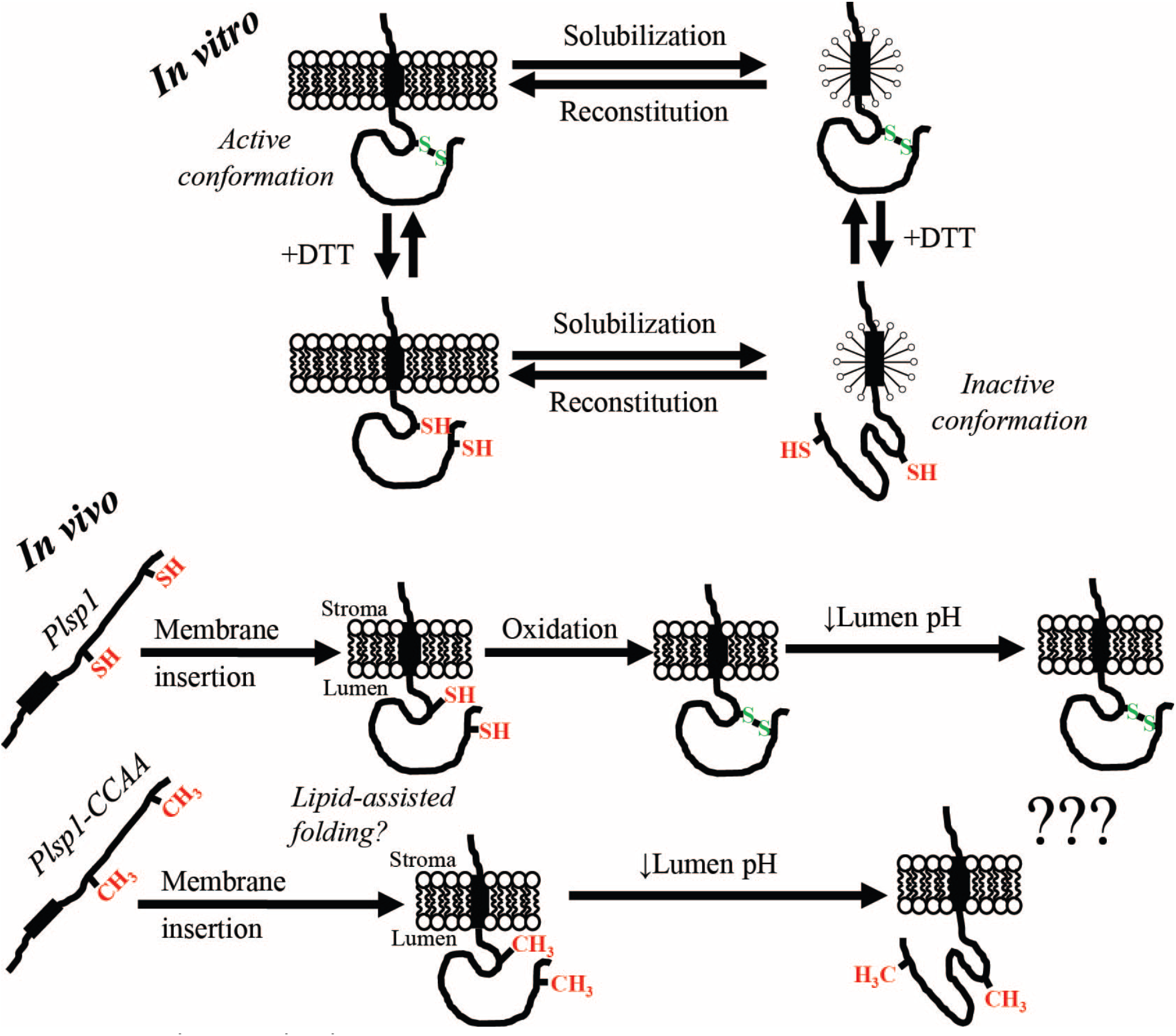
*In vitro* and *in vivo* models. *(In vitro)* Plsp1 is solubilized in from membranes in an active form due to an intramolecular disulfide bond. Reduction of this disulfide bond causes an inactivating conformational change in detergent micelles that is prevented by association with a lipid bilayer. Reconstitution of Plsp1 into a bilayer causes refolding back into the active state. **(*In vivo*)** Plsp1 is targeted to thylakoids in a reduced form. Once the catalytic domain traverses the membrane, folding into the catalytically active form is assisted by bilayer lipids. In the case of the wild-type protein, oxidation to form the disulfide bond then takes place and stabilizes the structure. Structural stabilization by the disulfide bond allows Plsp1 to remain optimally active amidst fluctuations in the lumen pH.

How do lipids make Plsp1 insensitive to the effects of reducing the disulfide bond? The soluble catalytic domain of *E. coli* LepB spontaneously inserts into membranes via its proposed hydrophobic membrane association surface (Bhanu and Kendall, 2014; van Klompenburg et al., 1998). A similar hydrophobic surface is revealed in the predicted structure of Plsp1 and may be responsible for the propensity of Plsp1 to associate with isolated thylakoid membranes (Endow et al., 2015). It is possible that insertion of the catalytic domain of Plsp1 into a membrane rigidifies the tertiary structure and prevents inactivating conformational changes resulting from disulfide bond reduction. However, there are still likely to be some conformational differences between oxidized and reduced Plsp1 despite being integrated into a membrane since the protease susceptibility of the wild type and single Cys variant in PLs are different (Fig. 7B). Another possible explanation is related to signal peptide-binding. Type I signal peptidases possess a specific groove into which a signal peptide binds (Paetzel, 2014). Midorikawa et al (2014) hypothesized that the C-terminal region can sterically interfere with binding of a thylakoid transfer signal (TTS) when Plsp1 is in a reduced state. If this is true, perhaps the catalytic domain of Plsp1 inserts into the membrane such that the TTS-binding groove becomes inaccessible to the C-terminal segment. An analogous example of intramolecular inhibition can be seen in Rv1827 from *Mycobacterium tuberculosis* in which phosphorylation of an N-terminal Thr residue causes this region of the protein to occlude a surface involved in interactions with binding partners (Nott et al., 2009).

Our results clearly indicate that lipids play an important role in the folding and activity of Plsp1, but they do not reveal the mechanism by which a membrane facilitates a reversible transition from an inactive to a catalytically active conformation. In *E. coli*, the multi-pass lactose permease (LacY) requires interactions with phosphatidylethanolamine (PE) to attain proper membrane topology and folding of a specific periplasmic domain (Dowhan et al., 2004). Once LacY is properly folded, interactions with PE are no longer required (Bogdanov and Dowhan, 1999). The mechanism of lipid-assisted folding of Plsp1 is likely different since Plsp1 only has a single transmembrane alpha helix, and the stimulatory effects of lipids on the single Cys Plsp1 variant are reversed by solubilization with Triton X-100 (Fig. 7C and 7D). Thus, the interaction between lipids and Plsp1 in membranes is likely stable as opposed to the transient nature of the interaction between PE and LacY (Bogdanov and Dowhan, 1999). Perhaps the mechanism with Plsp1 is similar to that of the membrane insertion of the pore-forming domain of colicin A (Lakey et al., 1991). Specifically, interactions with negatively charged lipid head groups are proposed to induce rearrangements of several amphipathic helices leading to exposure of two hydrophobic helices and subsequent membrane insertion (Lakey et al., 1991). Another possibility, albeit completely speculative, is that the C-terminal region of Plsp1 occupies the TTS-binding groove when the disulfide bond is lacking, as suggested above, but is displaced by lipids during membrane insertion.

There are many examples of proteins that require specific lipid molecules for activity (Hansen et al., 2011; Latowski et al., 2000), structural stability (Seiwert et al., 2017), proper folding (Dowhan et al., 2004), or membrane insertion (Mel and Stroud, 1993). In some cases, this is due to properties of the lipid head group such as charge (Hansen et al., 2011; Mel and Stroud, 1993), and in others it is related to biophysical properties of the lipid (Latowski et al., 2004; Seiwert et al., 2017). Although our current data do not reveal an effect of a specific lipid molecule, we noted already that liposomes made from thylakoid lipids are significantly more effective in maintaining Plsp1 activity than are those made from *E. coli* lipids. The most abundant lipid found in thylakoid membranes (MGDG) is a non-bilayer forming lipid (Dormann and Benning, 2002) and is important for the structure and function of various thylakoid proteins (Latowski et al., 2000; Seiwert et al., 2017). MGDG is also the only non-bilayer forming lipid used in our liposomes. PE is a non-bilayer forming lipid that mediates insertion of *E. coli* LepB into membranes and is also the most abundant phospholipid in *E. coli* (van Klompenburg et al., 1998). It would be interesting to test whether non-bilayer forming lipids are important for the activity and redox-dependency of Plsp1 in PLs by replacing MGDG with PE or phosphatidylcholine (a bilayer-forming lipid) (Latowski et al., 2004) (van Klompenburg et al., 1998).

### Does oxidative folding stabilize Plsp1 during diurnal changes in the energetic state of the thylakoid membrane?

In liposome reconstitution experiments, we found that a single Cys Plsp1 mutant has comparable activity to the wild-type form at pH 8 but lower comparative activity at pH 5.6 (Fig. 7E and F). This suggests that the oxidized form of Plsp1 is more stable than the reduced form under moderately acidic conditions, despite the fact that the membrane alone is sufficient to facilitate proper folding. The lumen pH drops when thylakoids are energized and can fluctuate in response to changes in light intensity (Shikanai and Yamamoto, 2017). Therefore, we hypothesize that the disulfide bond helps maintain Plsp1 structure and activity in response to changes in lumen acidity (Fig. 8). By altering protonation states, changes in pH could disrupt intramolecular interactions or interactions between Plsp1 and lipid head groups that are important for folding.

It is interesting to consider whether this is a property specific to Plsp1 or whether it applies to other proteins in general. One thought is that it may apply more often in proteins found in energy-transducing membranes. Such membranes give rise to the possibility that the protein will be found in rather different environments depending on the state of energization, which itself depends on external factors, such as availability of substrates for oxidative electron transport and light for photosynthetic electron transport. Hamsanathan and Musser (2018) recently considered explicitly that the native environment of a protein embedded in an energetic membrane would necessarily include the pmf. Indeed, an influence of an electric field on protein stability has been observed experimentally (Bekard and Dunstan, 2014) and by simulations (Jiang et al., 2019). Extramembranous effects of the pmf necessarily include the pH values in the flanking compartments, i.e., the thylakoid lumen in the case of Plsp1. The importance of the membrane for the folding of Plsp1 is noteworthy given that the active site is most likely near the membrane/lumen interface (Dalbey et al., 2012).

Many enzymes have evolved to function in extreme environments such as high temperatures, low pH, and high salt concentrations (D’Amico et al., 2003; van den Burg, 2003). Under such circumstances, those proteins exhibit properties adapted to the particular conditions encountered, and in fact, those conditions must prevail for maximal activity. For example, the maximum *in vitro* activities of alpha amylases from a psychrophile and a thermophile were observed near the temperatures at which the host organisms are adapted to growing (D’Amico et al., 2003). This reflects the fact that the conditions under which the proteins are evolved to handle are relatively stable. This contrasts to the proteins residing in the energy-transducing membranes in chloroplasts and photosynthetic bacteria, which can exist in the presence of a variable pmf (i.e., magnitude and composition) resulting from rapidly changing light conditions during the day (Armbruster et al., 2017; Shikanai and Yamamoto, 2017) and darkness at night. Such proteins must be able to adapt to a more variable environment than those that exist under stable but harsh environments. The oxidative folding of Plsp1 may be an example of such an adaptation to a changing energy environment. Other examples may include STN7, another thylakoid membrane protein that is likely stabilized by a lumenal disulfide bond (Shapiguzov et al., 2016). While the disulfide bond in STN7 was proposed to maintain the conformation necessary for its interaction with the cytochrome b6f complex rather than simply enhancing protein stability (Shapiguzov et al., 2016), this was not investigated under energizing and non-energizing conditions.

A puzzling feature of oxidative folding of Plsp1 is the fact that it is not conserved throughout all photosynthetic organisms but is apparently restricted to the angiosperms (Midorikawa et al., 2014). Homologs of Plsp1 are found in the bacterial plasma membrane and the mitochondrial inner membrane and cleave signal sequences from proteins targeted to the periplasm and intermembrane space, respectively (Paetzel et al., 2002). In contrast to these energy-transducing membranes, thylakoids store a major proportion of their total pmf as a ΔpH (Cruz et al., 2001). While it is tempting to speculate that oxidative folding evolved in Plsp1 as a general adaptation to functioning in a compartment that undergoes more drastic pH changes, we recognize that lumen acidification is likely a feature of all organisms that perform oxygenic photosynthesis, not just the angiosperms. One possibility is that all non-angiosperm Plsp1 orthologs possess a mechanism other than oxidative folding to stabilize the structure amidst pH changes. Another possibility is that conservation of Cys in angiosperm Plsp1 orthologs is related to the differences in pmf regulation between angiosperms and other members of the green lineage (Shikanai and Yamamoto, 2017). While the pmf composition can vary depending on environmental conditions (Shikanai and Yamamoto, 2017), perhaps the contribution of ΔpH is more variable in angiosperms compared to other photosynthetic organisms.

### Conclusion

The role of lipids in protein structure and function has gained increasing attention in recent decades. An important role of lipids in folding and structural stability could be a feature of many integral and peripheral proteins in all membranes, but structural stabilization by disulfide bonds may be most common in thylakoid membranes. This may be due to the fact that the thylakoid is the only major energy-transducing membrane which supports a substantial and highly dynamic pH gradient. Additionally, the apparent lack of major chaperones in the thylakoid lumen (Kieselbach and Schroder, 2003; Peltier et al., 2002; Schubert et al., 2002) may necessitate other means to stabilize proteins under periods of stress. While not discussed here, the electrical component of the pmf can also impact protein functions (Bekard and Dunstan, 2014; Jiang et al., 2019). Overall, we suggest our work has the potential to prompt and inform future discussions about the role of lipids and electrochemical gradients in protein structure and function.

## Materials and Methods

### Antibodies

The Pea Plsp1 antibody was from crude antiserum described previously (Midorikawa et al., 2014). The antibody against the extreme C-terminus of Arabidopsis Plsp1 was as described (Shipman and Inoue, 2009). The antibodies against OE23 and Toc75 were as described previously (Shipman-Roston et al., 2010), and that against Hcf106 was as described (Rodrigues et al., 2011). The OE33 antibody was a gift from Dr. Hsou-Min Li (Institute of Molecular Biology, Academia Sinica), that against PGRL1 was from AgriSera (Vännäs, Sweden), that against GFP was from Santa Cruz Biotech (https://www.scbt.com), and the polyclonal T7 antibody was obtained from Millipore (http://www.emdmillipore.com/US/en). Western blots were developed either by enhanced chemiluminescence using Luminata^TM^ Crescendo Western HRP Substrate or colorimetrically using the reagents 5-bromo-4-chloro-3-indolyl phosphate and nitroblue tetrazolium.

### Plant materials

The *pgrl1ab* double mutant was generated by a cross between SAIL_443E10 and SALK_059233 (DalCorso et al., 2008). Arabidopsis seeds, ecotype Columbia-0 and the *plsp1-1* T-DNA mutant (SALK_106199), were obtained from the Arabidopsis Biological Resource Center (https://abrc.osu.edu/).

### DNA constructs

The pUNI51 plasmid containing the coding sequence of Arabidopsis PsbP1 (At1g06680) was obtained from the Arabidopsis Biological Resource Center. The plasmid containing the coding sequence of the *Silene pratensis* plastocyanin precursor protein was as described (Last and Gray, 1989).

The cDNAs encoding Plsp1 variants for expression *in planta* were prepared by long flanking homology PCR as described previously (Endow and Inoue, 2013) using the primers listed in table S1 with *PLSP1*_*1-70*_-*CITRINE-PLSP*_*171-291*_ in pMDC32 as the template. PCR products were recombined into pDONR207 by a BP reaction using Gateway™ BP Clonase™ II Enzyme mix (Invitrogen). Gateway™ LR Clonase™ II Enzyme mix (Invitrogen) was then used for recombination into pMDC32. Each plasmid was confirmed by DNA sequencing and transformed into *Agrobacterium* GV3101 cells for plant transformation.

### Transient expression in Nicotiana benthamiana

*Agrobacterium* cells carrying the P19 suppressor (Voinnet et al., 2003) in pBIN69 or *PLSP1*_*1-70*_-*T7-PLSP171-291* in pMDC32 were grown overnight at ~28°C in LB broth with Kanamycin (25 μg/mL), Rifampicin (35 μg/mL) and Gentamycin (25 μg/mL). Cell cultures were then diluted in LB with Kanamycin (25 μg/mL) and grown to an OD_600_ of ~0.2. Cells were pelleted at 2500 x *g* and resuspended in 10 mM MES-KOH pH 5.6, 1 mM MgCl_2_, 0.2% w/v glucose, 150 μM acetosyringone (induction medium) to an OD_600_ of ~0.4. T7-Plsp1 and P19 cell cultures were mixed at a ratio of 4:1 and incubated with moderate shaking at ~28°C for 2 hours. Cells were pelleted again at 2500 x *g* and resuspended in 5% w/v sucrose, 300 μM acetosyringone (infiltration medium). *Nicotiana benthamiana* leaves were co-infiltrated using a syringe. After leaf infiltration, the plants were held in dark overnight and returned to a growth chamber (12 hours light per day, 20°C) for 1-3 days before isolating chloroplasts or thylakoids.

### Chloroplast isolations

*Nicotiana benthamiana* chloroplasts were isolated from leaves 2-4 days after infiltration with *Agrobacterium* cells. Leaves were homogenized in At grinding buffer (50 mM HEPES-KOH pH 8, 0.33 M sorbitol, 2 mM EDTA, and 0.5% w/v BSA) and filtered through two layers of Miracloth (Millipore). Pelleted chloroplasts (3000 x *g*, 4 °C, 3 minutes) were resuspended in a small volume of At grinding buffer and were carefully layered on top of a 40%/80% Percoll (GE Healthcare) step gradient (24 mL 40% Percoll on top of 6 mL 80% Percoll, both in At grinding buffer). After centrifugation at 4000 x *g* for 10 minutes, the green band at the interface (intact chloroplasts) was collected and diluted with ~25 mL of import buffer (50 mM HEPES-KOH or 50 mM Tricine-KOH, 0.33 M sorbitol, pH 8). After centrifugation at 2600 x *g* for 5 minutes, the intact chloroplasts were resuspended in import buffer, and a small volume was used for chlorophyll quantification as described (Arnon, 1949). The remainder was centrifuged again at 2600 x *g* for 5 minutes, and the pellet was resuspended in import buffer to 1 mg Chl/mL.

*Arabidopsis thaliana* chloroplasts were isolated from plants grown on phytoagar plates containing Murashige-Skoog with Gamborg’s vitamins (Caisson Laboratories), supplemented with 1% w/v sucrose, in a manner similar to that for *N. benthamiana* chloroplast isolation. After filtration through miracloth and centrifugation, chloroplasts were resuspended in At grinding buffer and layered on top of a 50% continuous Percoll gradient. The green band near the bottom of the gradient after centrifugation at 8000 x *g* for 10 minutes was collected into ~25 mL of import buffer, and chloroplasts were pelleted and washed in import buffer as above. Chlorophyll quantification and resuspension to 1 mg Chl/mL were carried out as described above.

Chloroplasts from *Pisum sativum* plants (Little Marvel) were isolated from 11-14 day-old plants exactly as was done for Arabidopsis chloroplasts with the following exception: Pea grinding buffer (50 mM HEPES-KOH pH 8, 0.33 M sorbitol, 2 mM EDTA, 1 mM MgCl_2_, 1 mM MnCl_2_, and 0.1% w/v BSA) was used in place of At grinding buffer.

### Protease treatments of thylakoid extracts

Pea chloroplasts isolated as described above were lysed hypotonically at ~1 mg Chl/mL in 1 mM Tricine-NaOH pH 7, 5 mM MgCl_2_ (Buffer A) for 10 minutes on ice in the dark. Membranes were pelleted at 5000 x *g* for 5 minutes and were washed three times with 10 mM Tricine-NaOH pH 7, 5 mM MgCl_2_ 0.3 M sucrose (Buffer B). Washed thylakoids were resuspended in buffer C (50 mM Tricine-NaOH pH 7, 5 mM MgCl_2_, 15 mM NaCl) to ~2 mg Chl/mL. An equal volume of buffer containing 0.5% v/v Triton X-100 was added, and the sample was incubated for 10 minutes on ice in the dark. After centrifugation at 100,000 x *g*, 4°C, 8 minutes (TLS55 rotor), the light-green supernatant (thylakoid extract) was transferred to a new tube and used immediately for experiments. Thylakoid extracts were pre-treated with or without 10 mM TCEP (Thermoscientific) for 20 minutes on ice in the dark. Aliquots of each were then mixed with buffer containing CaCl_2_ (0.5 mM final) with or without thermolysin (0.1 μg/mL final). Each reaction mixture was incubated at 25°C. Aliquots taken at 10, 20, and 30 minutes were mixed with an equal volume of 2X reducing (0.2 M β-ME) sample buffer containing 20 mM EDTA and were boiled for 5 minutes. Samples were then analyzed by SDS-PAGE and immunoblotting or Coomasie Brilliant Blue staining.

### Affinity purification of T7-Plsp1

T7-Plsp_171-291_ (T7-Plsp1) was extracted from *N. benthamiana* thylakoids after transient expression of *PLSP1*_*1-70*_-*T7-PLSP*_*171-291*_ in leaves. Leaves were homogenized in Pea grinding buffer with 244 μM PMSF, filtered through four layers of Miracloth, and the filtrate was centrifuged at 2500 x *g* for 4 minutes at 4°C. The resulting pellet was washed once with 50 mM Tricine-NaOH pH 7, 5 mM sorbitol and three times with 50 mM Tricine-NaOH pH 7, 5 mM MgCl_2_, 0.1 M sorbitol. For extraction with Triton X-100, crude thylakoids were resuspended in buffer C (see above) at ~2 mg Chl/mL and mixed with an equal volume of the same buffer containing 0.5% v/v Triton X-100. For extraction with octyl glucoside, crude thylakoids were resuspended in buffer C at ~1 mg Chl/mL, and solid octyl glucoside was added to 1% w/v. After mixing at 4°C in the dark for 10-30 minutes, the supernatant containing T7-Plsp1 was obtained by centrifugation at 30,000 x *g* for 30 minutes at 4°C. T7-Plsp1 was then bound to T7•Tag Antibody Agarose (Millipore) for 1-2 hours at 4°C on a BioRad EconoPac 25mL column. The column was washed once with buffer C containing the appropriate detergent (0.25% v/v Triton X-100 or 1% w/v octyl glucoside) and was eluted with 0.1 M Glycine-HCl pH 2.2 containing the same detergent. Elution fractions were collected into 4 M Tris-HCl pH 9.5 (20 μL/mL elution) to give a final pH of ~7.5. Each fraction was analyzed by SDS-PAGE and Coomasie Brilliant Blue staining or immunoblotting using the indicated antibodies. Purified T7-Plsp1 was divided into numerous aliquots which were frozen with liquid N_2_ and stored at −80°C.

### Synthesis of radiolabeled proteins

Proteins were synthesized from plasmid DNA templates in the presence of ^35^S (EasyTag™ EXPRESS35S Protein Labeling Mix, Perkin Elmer) using rabbit reticulocyte lysate (TnT® Quick Coupled Transcription/Translation System, Promega) according to the manufacturer’s guidelines. Reaction mixtures were quenched with an equal volume of 50 mM L-Methionine, 15 mM L-Cysteine in 0.1 M HEPES-KOH pH 8, 0.66 M sorbitol and were used for experiments on the same day.

### Fluorescence analysis of Plsp1

Aliquots of purified T7-Plsp1 in 1% w/v octyl glucoside were thawed on ice. Emission spectra of 150 μL samples of T7-Plsp1, T7-Plsp1 with 50 mM DTT, and the corresponding buffer blanks all prepared in elution buffer neutralized with Tris-HCl were collected using a Fluorolog spectrofluorometer (Horiba Scientific). The final concentration of T7-Plsp1 in each sample was ~0.1 μM as determined by comparison to a BSA standard curve on a Coomasie-stained SDS-PAGE gel. Samples were excited at 285 nm, and slit widths were set to 5nm for both excitation and emission. T7-Plsp1 with 50 mM DTT was incubated on ice for 20 minutes prior to collecting the emission spectrum. Spectra of buffer blanks were subtracted from those of each T7-Plsp1 sample to give corrected emission spectra.

### Thiol labeling assay

Intact chloroplasts were isolated from *Nicotiana benthamiana* leaves after *Agrobacterium*-mediated transient expression of Plsp1_1-70_-T7-Plsp_171-291_ as described above or after infiltration with infiltration medium (mock). Chloroplasts from mock-infiltrated leaves were subject to mock treatments at each step. Chloroplasts containing T7-Plsp1 were hypotonically lysed in the presence or absence of 0.1 M NEM and washed two times. Washed chloroplast membranes were then solubilized for 30 minutes at 28°C with 0.2 M Bis-Tris-HCl pH 6.5, 2% w/v SDS (labeling buffer) with or without 0.1 M NEM. Proteins were precipitated with an equal volume of 20% TCA/80% acetone, and pellets were washed 3-4 times with 1 mL 80% acetone. Protein pellets were then resuspended in labeling buffer with or without 2 mM TCEP and incubated for 20 minutes at 28°C. Labeling buffer with or without methoxypolyethylene glycol maleimide (mPEG-MAL, Sigma) was then added to 10 mM final, and samples were then incubated for another 2 hours at 28°C. Proteins were again precipitated as above and were resuspended in 2X sample buffer (0.1 M Tris-HCl pH 6.8, 4% SDS, 20% glycerol, 0.2% bromophenol blue, 0.2 M β-ME). Sample were analyzed by SDS-PAGE and immunoblotting.

### Generation and identification of transgenic Arabidopsis plants

Stable transgenic Arabidopsis plants were obtained by floral dip transformation of heterozygotes for the *plsp1-1* T-DNA insertion using *Agrobacterium tumefaciens* cell cultures carrying the indicated transgene in pMDC32. T1 seeds were surface sterilized with 35% bleach, 0.02% v/v Triton X-100 and then screened on MS medium containing 25 or 50 μg/mL hygromycin (Calbiochem). Hygromycin-resistant seedlings (i.e., visible true leaves, roots penetrating the agar medium, wild type-like appearance) were transferred to soil after 2-3 weeks and were grown at 16 hours light per day, 20°C. Soil-grown plants were screened by PCR using whole leaf DNA extracts. Selected plants were propagated to the T3 or T4 generation and used for experiments.

### Arabidopsis in vitro import assay

Intact chloroplasts were isolated as described above from 21-24 day-old Arabidopsis plants grown on MS agar medium at ~20°C under ~60 μE light with a 12 hour light per day regime. 50 μL import reactions containing 3mM MgATP, 10 μL radiolabeled precursor proteins, and chloroplasts at 0.22 mg Chl/mL were prepared on ice in the dark. Upon addition of the radiolabeled proteins, each sample was immediately moved to ambient light (10-30μE) at 20-22°C and incubated for 5, 10, or 20 minutes. Start times were staggered such that all import reactions would end at the same time. Samples were then immediately moved to ice in the dark and layered on top of 35% Percoll cushions in import buffer to re-isolate intact chloroplasts.

After centrifugation at 900 x *g*, 5 minutes, 4°C, all but ~50 μL of each supernatant was removed. 100 μL of import buffer was added, and the green pellets were resuspended by pipetting. 100 μL of each sample was then transferred to a new tube and centrifuged again at ~2000 x *g*, 5 minutes, 4°C. After discarding the supernatants, each green pellet was resuspended in 2X sample buffer and heated at ~82°C for 5 minutes. The leftover ~50 μL solutions were used for chlorophyll quantification. Samples were analyzed by SDS-PAGE and autoradiography with the loading normalized based on total chlorophyll. Radioactive bands were quantified using ImageJ 1.5i (National Institutes of Health). Signals were normalized to that of the Citrine-Plsp1 wild type sample at 20 minutes.

### Total protein extraction

Whole seedlings were frozen and homogenized under liquid N_2_ followed by suspension and boiling for ~15 minutes in extraction buffer (0.1 M Tris-HCl pH 6.8, 4% SDS, 15% glycerol, 10 mM EDTA, 2% β-ME). After centrifugation to remove tissue debris, four volumes of 100% acetone were added to the soluble extract for precipitation at −20°C. Precipitated proteins were collected by centrifugation, washed with 80% acetone, and resuspended in 8 M urea buffered with 0.1 M sodium phosphate and 10 mM Tris-HCl, pH 8.6. Total proteins were quantified by the Bradford assay (Bradford, 1976) using BSA as the standard.

### In vitro processing assays

A typical 10 μL processing assay consisted of 1 μL radiolabeled protein, enzyme sample (purified Plsp1 or crude thylakoid extract), and enzyme buffer with or without various additions such as DTT. Pre-incubations without substrate were typically carried out on ice for ~30 minutes. Reaction mixtures were incubated at 25-30°C for 30 minutes and quenched by addition of 10 μL of 2X sample buffer. After boiling for 5 minutes, samples were analyzed by SDS-PAGE and autoradiography.

Permeablization of thylakoid membranes was carried out as described previously (Ettinger and Theg, 1991). Washed thylakoids were prepared as done for protease treatment experiments described above. Samples at ~1 mg Chl/mL were mixed with an equal volume of buffer C (see above) containing 0.08% v/v Triton X-100 and incubated on ice in the dark for 10 minutes. Thylakoid extracts were prepared as described above. For processing assays using permeablized thylakoids (A) or thylakoid extracts (B), 8 μL of enzyme sample was mixed with 1 μL of buffer containing 0.08% v/v Triton X-100 (A) or 0.5% v/v Triton X-100 (B) and 1 μL of radiolabeled substrate protein. For pre-treatments, the buffer contained the indicated chemical at a 10X concentration. Samples were incubated at ~25°C for 30 mins in the dark.

### Reconstitution of T7-Plsp1 into liposomes

Purified T7-Plsp1 in 0.25% v/v Triton X-100 was reconstituted into liposomes composed of *E. coli* total lipid extract or a mixture of thylakoid lipids (19.8 mol% MGDG, 54.9 mol% DGDG, 14.8 mol% SQDG, 10.5 mol% PG). All lipids were from Avanti Polar Lipids. Small unilamellar vesicles (SUVs) were prepared by resuspending dried lipids at 4 mg/mL in liposome buffer (50 mM Tris-HCl pH 8, 50 mM KCl, 10% glycerol) and sonicating with a probe sonicator until optical clarity was achieved. Metal fragments from the sonicator tip were removed by centrifugation at 17,000 x *g* for 2 minutes. SUVs were mixed with Triton X-100 at a mass ratio of 2:1 to saturate the vesicles with detergent (Rigaud and Levy, 2003). This mixture was incubated on ice for 15 minutes before adding purified T7-Plsp1 in 0.25% v/v Triton X-100 at a lipid to protein mass ratio of ~80:1. After mixing at 4°C for 30 minutes, the mixture was added to an amount of BioBeads SM-2 (BioRad) equal to approximately ten times the mass of Triton X-100 present. The supernatant was added to a second batch of BioBeads after 1 hour of mixing at 4°C and to a third batch after 2 hours of mixing at 4°C. The third BioBeads incubation was done at 4°C for 18-20 hours. BioBeads were removed using a Promega spin column, and the flow through was centrifuged at 250,000 x *g* for 30 minutes at 4°C. The transparent pellet was resuspended in 100 μL of liposome buffer and then centrifuged at 16,000 x *g* for 2 minutes to remove any aggregates. The supernatant was used for *in vitro* processing assays, thermolysin treatments, or a carbonate wash (0.1 M Na_2_CO_3_).

For reconstitution experiments described in Figure S9, Plsp1 in 1% octyl glucoside was reduced with 50 mM DTT, and 50 mM DTT was present throughout the reconstitution procedure to maintain Plsp1 in a reduced state. PLs were pelleted as described above and resuspended in 70 μL of liposome buffer followed by centrifugation at 16,000 x *g* for 2 minutes to remove any aggregates. The supernatant was used for *in vitro* processing assays and thermolysin treatments. To confirm the redox state of Plsp1 before and after reconstitution, an aliquot was precipitated with 10% trichloroacetic acid followed by incubation in SDS-PAGE sample buffer (without β-ME) containing the thiol-specific reagent 4-Acetamido-4’-Maleimidylstilbene-2,2’-Disulfonic Acid (AMS) at a concentration of 10 mM.

## Abbreviations

Δψ: transmembrane electrical gradient
ΔpH: transmembrane pH gradient
DGDG: digalactosyl diacyl glycerol
DTT: dithiothreitol
LepB: leader peptidase
MGDG: monogalactosyl diacyl glycerol
PG: phosphatidyl glycerol
Plsp1: lastidic type I signal peptidase 1
pmf: proton motive force
PLs: proteoliposomes
SQDG: sulfoquinovosyl diacyl glycerol
TTS: thylakoid transfer signal
TPP: thylakoidal processing peptidase

## Acknowledgements

This work was supported by the Office of Basic Energy Sciences of the U.S. Department of Energy (grant DE-SC0017035 to S.M.T. and grant DE-FG02-08ER15963 to K.I.), University of California Davis Department of Plant Sciences Graduate Student Research Assistantship (to L.J.M.), and a Henry A. Jastro Graduate Research Award (to L.J.M.)

