## Supplementary data for "Lipid Chaperoning of a Thylakoid Protease Whose Stability is Modified by the Protonmotive Force"

### Supplementary data and figure legends

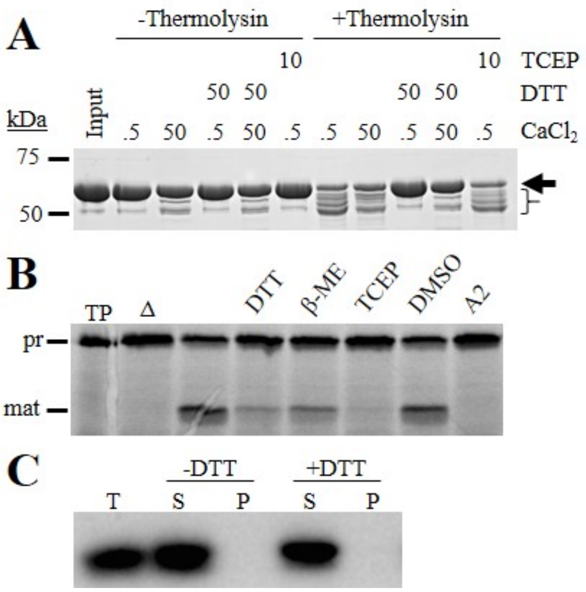


**Figure S1: Justification for using TCEP as a reductant, and Plsp1 solubility +/- DTT**

**(A)** 3 μg of recombinant Cpn60α was pre-treated with DTT or TCEP at the indicated concentration (in mM) or with buffer on ice for 25 minutes. Buffer containing $CaCl_2$ at the indicated concentration (in mM) with or without thermolysin (0.1 μg/mL final) was then added, and samples were incubated at 25°C for 15 minutes before analysis by SDS-PAGE and Coomasie staining. The arrowhead and brackets denote Cpn60α and degradation products, respectively. **(B)** Processing activity of purified T7-Plsp1 against prPsbP after treatment with various reducing agents. Purified T7-Plsp1 was pre-treated on ice for 30 minutes with or without 10 mM of the indicated reducing agent, 1% DMSO, or 10 μM Arylomycin A2 (A2) in 1% DMSO. After adding $^{35}$S-Met-prPsbP and incubating at 35°C for 30 minutes, an equal volume of 2X sample loading buffer was added. Samples were then boiled for 5 minutes and analyzed by SDS-PAGE and autoradiography. Δ = enzyme boiled for 10 mins prior to adding substrate. TP= translation products. **(C)** Purified T7-Plsp1 was incubated at 25°C in the presence (+) or absence (-) of 50 mM DTT for 30 minutes followed by centrifugation at 16,000 x *g* for 30 minutes at 4°C. The supernatant (S) was transferred to a new tube and mixed with an equal volume of sample loading buffer (SLB), and SLB was added the empty tube to resuspend any pelleted protein (P). Samples were analyzed by SDS-PAGE and immunoblotting using the antibody against Arabidopsis Plsp1.

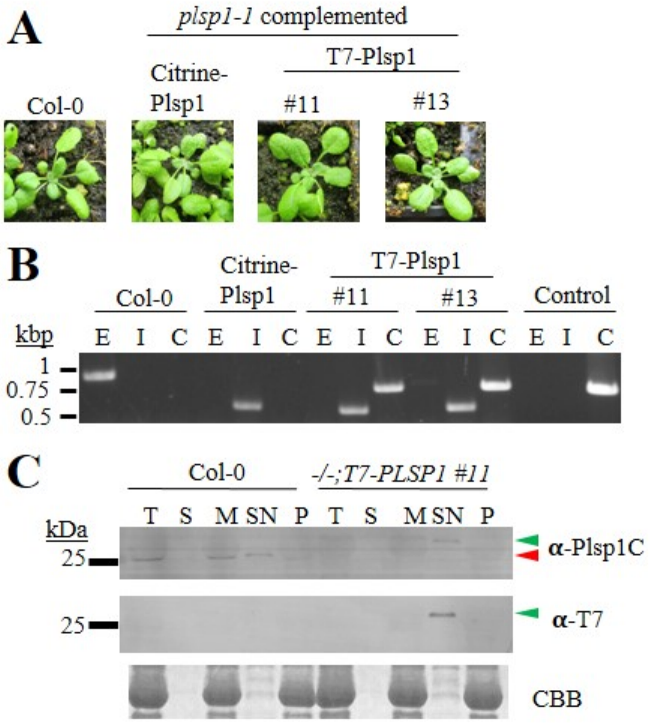


**<u>Figure S2. T7-Plsp1 is functional *in vivo*</u>**

**(A)** Images of 22 day-old soil-grown seedlings. **(B)** Genomic PCR results to confirm genotypes of seedlings shown in A. E = PLSP1, I = *plsp1-1* T-DNA insertion, C = tpPLSP1-T7-mPLSP1, Control= tpPLSP1-T7-mPLSP1 in pMDC32. **(C)** SDS-PAGE analysis of chloroplasts isolated from Col-0 wild type and T7-Plsp1-complemented seedlings. Green and red arrows denote T7-Plsp1 and Plsp1, respectively. The bottom panel shows the LHCP bands on a Coomasie-stained gel. T = total chloroplasts, S = chloroplast soluble fraction, M = membranes, SN and P = supernatants and pellets, respectively, after 0.25% Triton X-100 treatment.

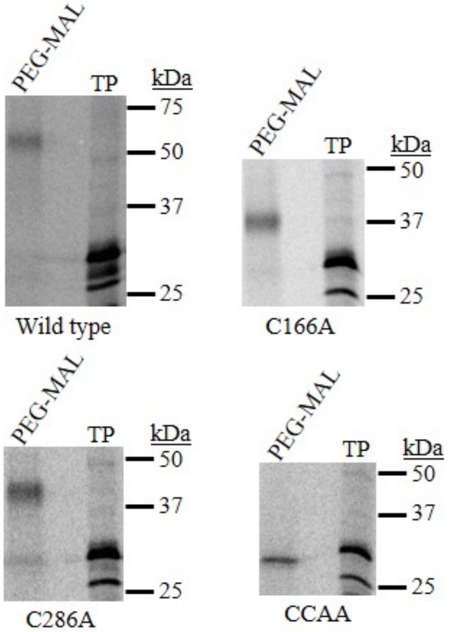


**Figure S3. PEG-MAL labeling of *in vitro* translated 10His-Plsp1$_{68\text{-}291}$**

*In vitro* translated 10His-Plsp1$_{68\text{-}291}$ variants labeled with $^{35}$S-Met were transported into Pea thylakoid membranes as described (Endow et al., 2015) but without thermolysin treatment. An aliquot of each post-transport mixture was treated with 5 mM TCEP in 50 mM Tris-HCl pH 7.8, 2% w/v SDS followed by addition of mPEG-MAL to 10 mM final. Samples were analyzed by SDS-PAGE and autoradiography. TP = translation products.

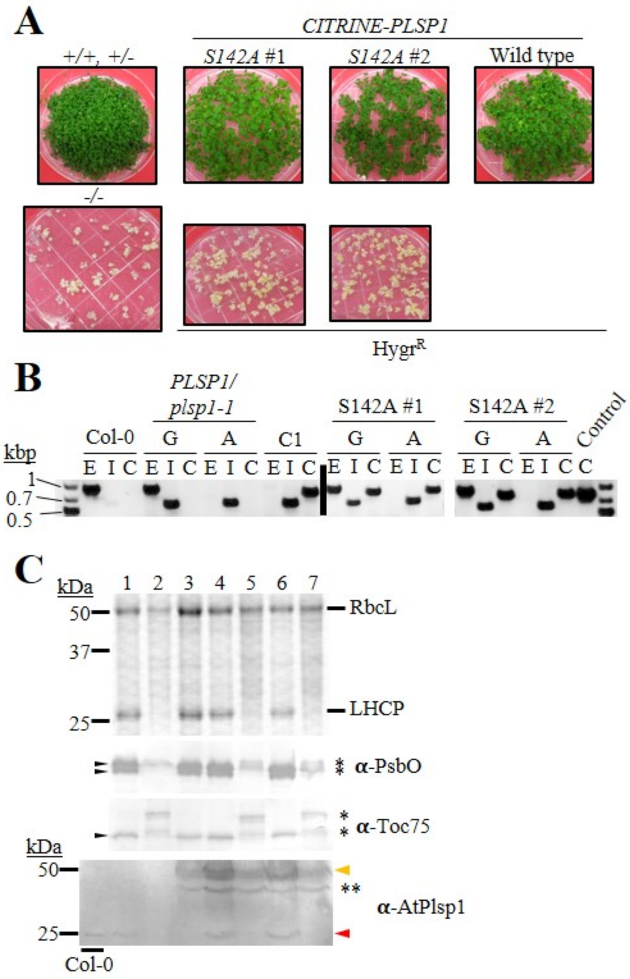


**<u>Figure S4. Plsp1 requires the catalytic nucleophile Ser142 for *in vivo* functionality</u>**

**(A)** Images of seedlings used for total protein extraction. Seeds were sown on MS medium supplemented with 3% w/v sucrose (*PLSP1/plsp1-1* segregating seeds) or 3% w/v sucrose with hygromycin at 25 µg/mL (*CITRINE-PLSP1* transgenic lines). Green seedlings from the *PLSP1/plsp1-1* line and hygromycin-resistant seedlings from the *CITRINE-PLSP1* transgenic lines were transferred to new MS medium containing 3% w/v sucrose without hygromycin after 7 and 14 days, respectively. Total proteins were extracted when seedlings were 31 (green) or 43 (albino) days old. (+/+, +/-) = mixture or homozygous wild type and heterozygous plants. (-/-) = *plsp1-1* null mutant. **(B)** Results of genomic PCR to confirm the genotypes of seedlings grown on MS medium. The vertical black bar indicates non-adjacent lanes of the same gel. G = green seedlings, A = albino seedlings, E = *PLSP1*, I = *plsp1-1*, C = *CITRINE-PLSP1*, asterisk indicates *CITRINE-PLSP1* in pMDC32. **(C)** Total protein extracts were analyzed by SDS-PAGE and Coomasie staining (top panel) or by immunoblotting (bottom panels) with the indicated antibodies. (1) +/+ and +/- plants, (Boomer, #792) *plsp1-1* null, (3) *CITRINE-PLSP1*-complemented, (4) +/+ or +/-;*CITRINE-PLSP1-S142A* #1, (5) -/-;*CITRINE-PLSP1-S142A* #1, (6) +/+ or +/-;*CITRINE-PLSP1*-S142A #2, (7) -/-;*CITRINE-PLSP1-S142A* #2. Gels were loaded with 210 ug BSA equivalents (α-AtPlsp1C) or 15 ug BSA equivalents (all others). Col-0 chloroplasts were used as a control for the α-AtPlsp1 immunoblot. The orange arrow indicates

Citrine-Plsp1, and the red arrow indicates Plsp1. The double asterisk indicates what may be a non-specific band or a Citrine-Plsp1 degradation product.

**A**

| Line | Sample sizes | % Hygr$^R$ | % Hygr$^S$ |
|---|---|---|---|
| C1 | 122, 82, 96 | 99.6 ± 0.6 | 0.3 ± 0.6 |
| C2* | 89, 101, 98 | 77.8 ± 4.7 | 22.2 ± 4.7 |
| C3* | 99, 104, 82 | 78.8 ± 11.7 | 21.2 ± 11.7 |
| CA1-1 | 98, 109, 74 | 100 | 0 |
| CA1-2* | 96, 109, 81 | 74.9 ± 1.4 | 25.1 ± 1.4 |
| CA1-3* | 85, 99, 82 | 77.0 ± 1.5 | 23.0 ± 1.5 |
| CA2-1 | 96, 114, 85 | 100 | 0 |
| CA2-2 | 88, 106, 88 | 99.3 ± 0.6 | 0.7 ± 0.6 |
| CA2-3 | 104, 101, 85 | 99.3 ± 0.6 | 0.7 ± 0.6 |
| CCAA1* | 90, 115, 87 | 79.3 ± 3.3 | 20.7 ± 3.3 |
| CCAA2 | 89, 102, 83 | 99.6 ± 0.6 | 0.4 ± 0.6 |
| CCAA3 | 103, 104, 98 | 99.0 ± 1.0 | 1.0 ± 1.0 |

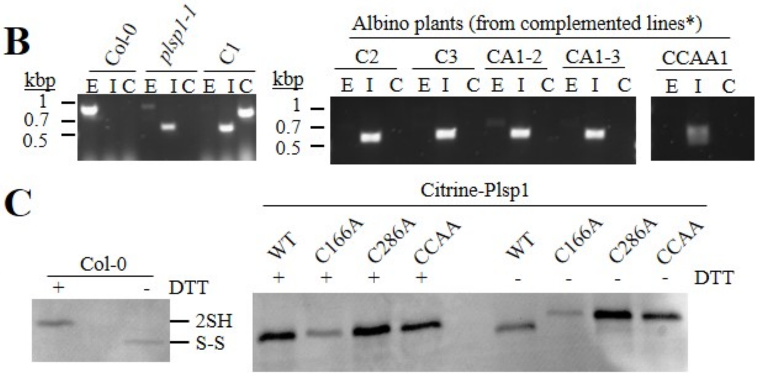


**Figure S5. Analysis of albino plants from hemizygous complemented lines and Citrine-Plsp1 mobility on SDS-PAGE**

**(A)** Seedlings grown on MS medium with 1% sucrose and hygromycin (25 µg/mL) were scored at 10-14 days-old. Hygromycin resistance was scored when seedlings exhibited visible true leaves, roots penetrating into the growth medium, and appeared green. Hygromycin susceptibility was scored when seedlings had no visible true leaves, appeared pale or albino, and had little or no visible root growth and penetration into growth medium. Asterisks indicate those lines that were hemizygous for the *CITRINE-PLSP1* transgene. **(B)** Genomic PCR results to confirm genotypes of the albino seedlings from lines C2, CA1-2, CA1-3, and CCAA1. PCR data in the left panel are identical to those shown in Figure 3 (upper panel). **(C)** Thylakoid extracts (0.25% v/v Triton X-100 supernatant) were treated with or without 50 mM DTT for 30 minutes on ice prior to non-reducing SDS-PAGE and immunoblotting using the α-AtPlsp1antibody. Samples from Col-0 (top panel) were run on a 12% gel, and all others (bottom panel) were run on a 7.5% gel to resolve oxidized and reduced Citrine-Plsp1. 2SH = reduced Plsp1. S-S = oxidized Plsp1.

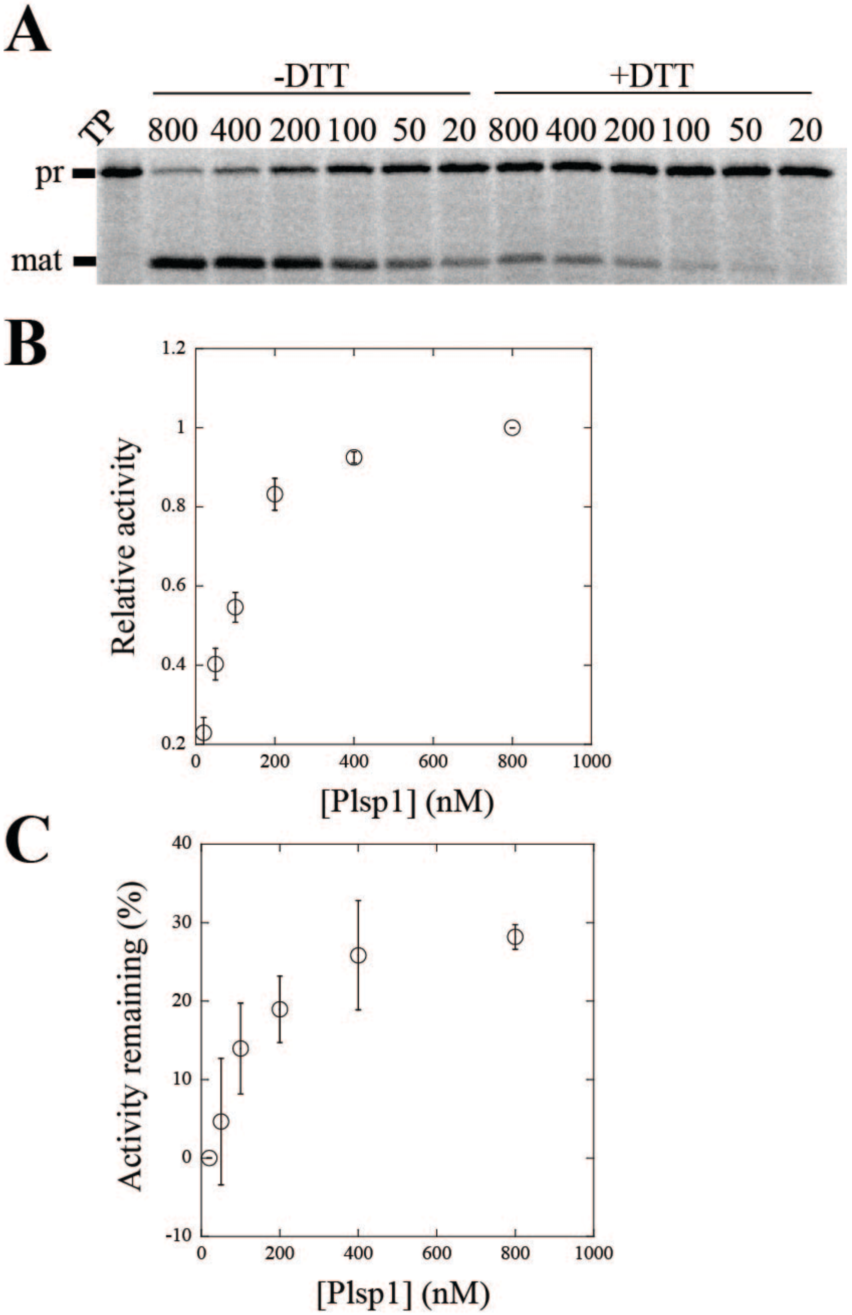


**Figure S6. The amount of inhibition of Plsp1 activity by DTT is dependent on the enzyme concentration**

**(A)** Purified T7-Plsp1 was incubated at various concentrations (in nM) indicated above each lane with or without 50 mM DTT on ice for 30 minutes. $^{35}$S-Met-prPsbP was then added to each enzyme sample followed by incubation at 25°C for 30 minutes. After adding an equal volume of 2X sample loading buffer and boiling for 5 minutes, samples were analyzed by SDS-PAGE and autoradiography. TP = translation product. **(B)** Quantification of activity of Plsp1 without DTT treatment versus Plsp1 concentration expressed as the mature PsbP signal relative to that for 800 nM Plsp1. Shown are the mean relative activities ± standard deviation of three independent experiments. **(C)** Quantification of effect of 50 mM DTT on Plsp1 activity expressed as the apparent % activity inhibited.

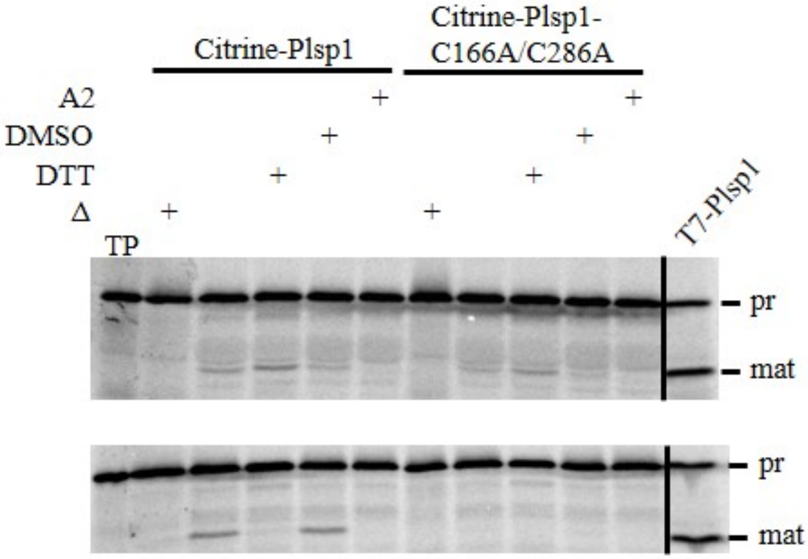


**Figure S7. Processing activity of Citrine-Plsp1-CA is lost upon extraction from thylakoids**
*In vitro* processing activity of 0.04% v/v Triton X-100-treated thylakoids (top panel) or of 0.25% v/v Triton X-100 thylakoid extracts (bottom panel) from Citrine-Plsp1 or Citrine-Plsp1-C166A/C286A-complemented Arabidopsis plants. Samples were pre-treated on ice for 30 mins with reaction buffer, 50 mM DTT, 1% v/v DMSO, or 10 µM Arylomycin A2 (A2) with 1% v/v DMSO. After adding $^{35}$S-Met-prPsbP, each reaction mixture was incubated 28°C for 30 minutes followed by addition of an equal volume of 2X sample loading buffer and boiling for 5 minutes. Samples were analyzed by SDS-PAGE and autoradiography. T7-Plsp1 was purified in 0.25% v/v Triton X-100 as described in Materials and Methods. TP = translation product. Δ = enzyme boiled for 5 minutes prior to adding substrate.

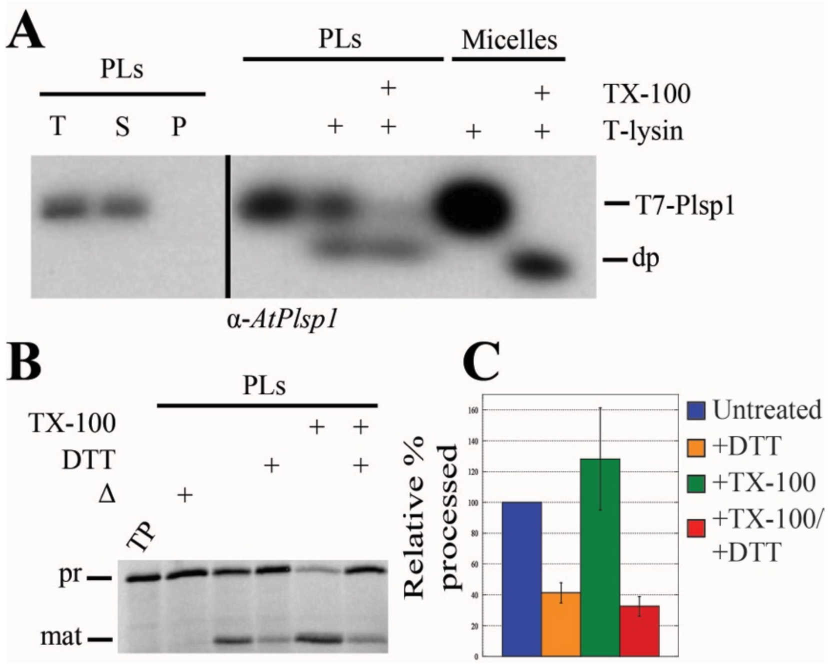


**<u>Figure S8. Reconstitution into *E. coli* lipid vesicles partially maintains Plsp1 activity under reducing conditions</u>**

**(A)** Purified T7-Plsp1 was reconstituted into liposomes made from *E. coli* total lipid extract as described in Materials in Methods. The recovered proteoliposomes (PLs) were treated with or without thermolysin (T-lysin) and/or 2% Triton X-100 (TX-100) and analyzed by SDS-PAGE and immunoblotting using the Arabidopsis Plsp1 antibody. T = total reconstitution mixture. S = soluble fraction after resuspending pelleted liposomes. P = pellet fraction after resuspending pelleted liposomes. **(B)** PLs were pre-treated with or without 50 mM DTT and/or 2% Triton X-100 (TX) followed by incubation with $^{35}$S-prPsbP as in Figure 7. Samples were analyzed by SDS-PAGE and autoradiography. Δ = boiled for 10 minutes prior to adding substrate. TP = translation product. **(C)** Quantification of mature PsbP bands in B relative to the untreated control. Shown are the means ± standard deviation from five independent experiments.

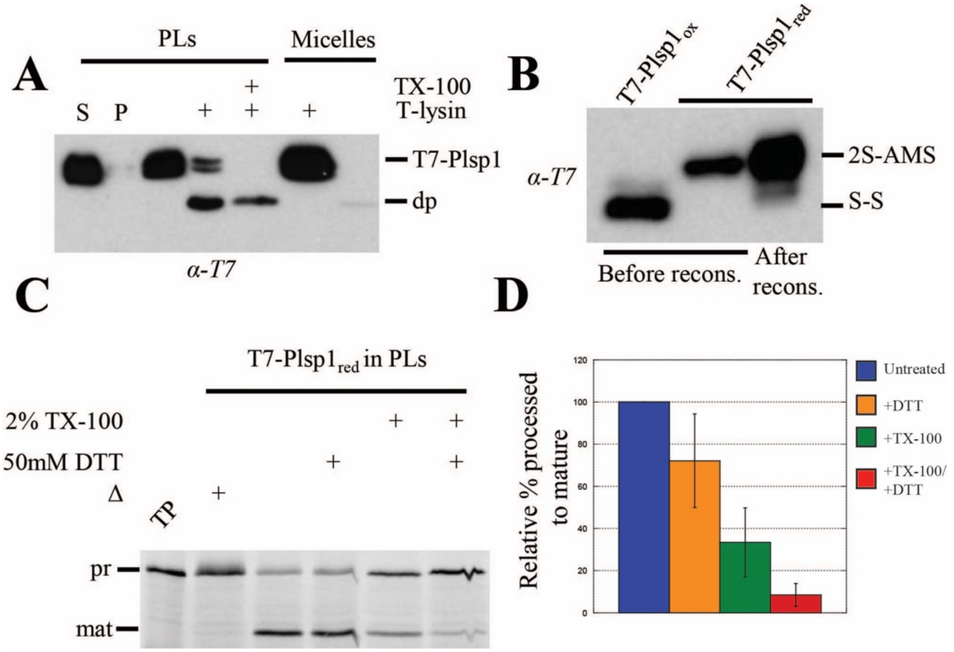


**Figure S9. Reconstitution of Plsp1 after reduction with DTT recovers processing activity**
**(A)** T7-Plsp1 in 1% octyl glucoside was treated with 50 mM DTT followed by reconstitution into liposomes made from thylakoid lipids as described in Materials and Methods except that 50 mM DTT was present throughout the reconstitution procedure. The recovered PLs were subjected to thermolysin treatment with or without 2% Triton X-100. S = soluble fraction after resuspending pelleted liposomes. P = pellet fraction after resuspending pelleted liposomes. **(B)** T7-Plsp1 in 1% octyl glucoside was treated with (red) or without (Dominguez Pardo, #1192) DTT followed by TCA precipitation and incubation in the presence of 10 mM AMS. An aliquot of the DTT-treated reconstituted T7-Plsp1 was also precipitated and treated with 10 mM AMS. 2S-AMS = reduced and AMS-labeled form of Plsp1. S-S = oxidized form of Plsp1. **(C)** PLs described in A and B were incubated with $^{35}$S-prPsbP as in Figure 7. Samples were analyzed by SDS-PAGE and autoradiography. **(D)** Quantification of mature PsbP bands in C relative to the untreated control. Shown are the means ± standard deviation from two independent reconstitution experiments.

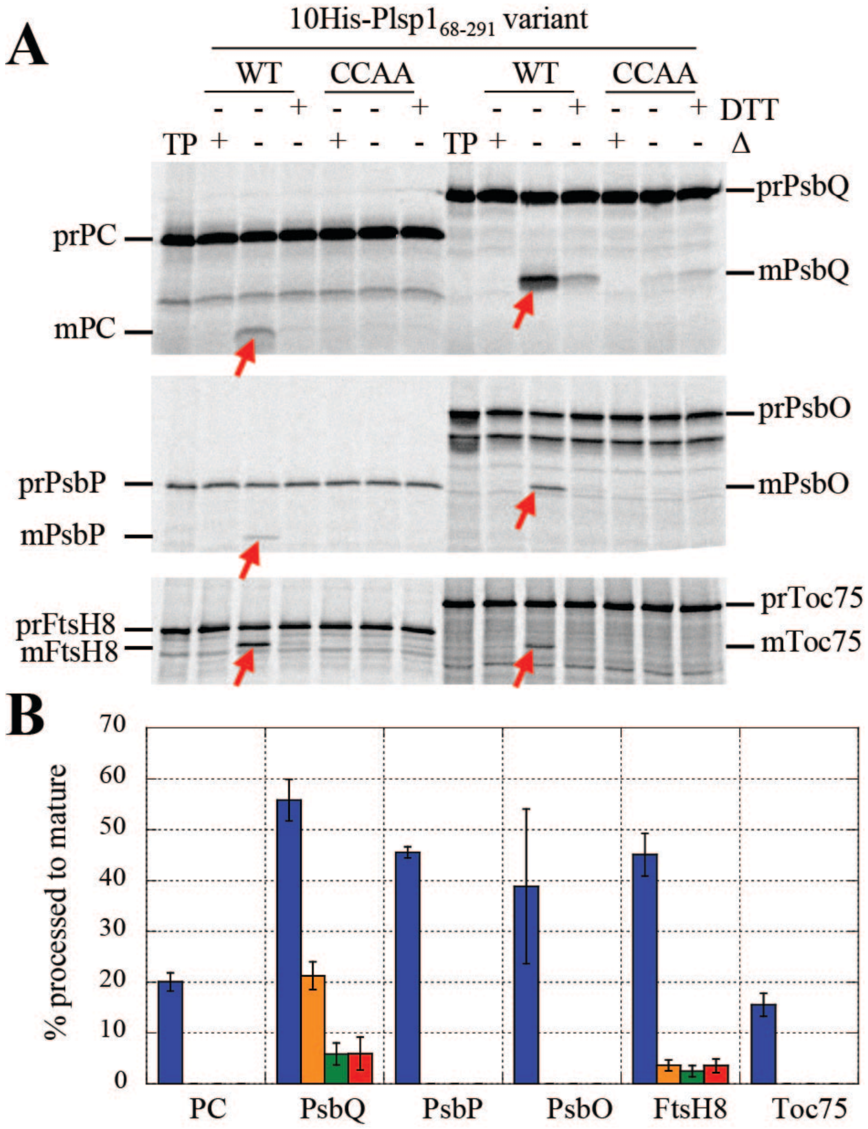


**Figure S10. Substituting both Cys in Plsp1 for Ala abolishes *in vitro* activity**

**(A)** Each 10His-Plsp1$_{68-291}$ variant was purified using NiNTA agarose and mixed with $^{35}$S-Met-labeled substrate and incubated at 25°C for 30 minutes. Samples were analyzed by SDS-PAGE and autoradiography. The final Plsp1 concentration was 100 nM in each processing reaction mixture. Red arrows indicate mature forms of each substrate tested. Substrates tested were precursor forms of plastocyanin (Zagari, #1096), PsbQ, PsbP, PsbO, FtsH8, and Toc75. TP = translation products. **(B)** Quantification of mature forms of each substrate in each sample. Data shown are expressed as the % of each precursor converted to the mature form (normalized to #Met residues in each form) and are means ± SD of three independent experiments. Wild-type Plsp1 without and with DTT are shown in blue and orange, respectively. The C166A/C286A mutant Plsp1 without and with DTT are shown, where applicable, in green and red, respectively. Where bars are absent is indicative of undetectable activity.

**List of primers used in this study**

| Name | Direction | Sequence (5'→3') | Purpose |
|---|---|---|---|
| PLSP1-F | Forward | AACGGATTGTTGCCAAAGAAGG | Genotyping |
| PLSP1-R | Reverse | GCAGCTTCCGACAAGAAGGGT | Genotyping |
| T-DNA left border | Reverse | ATTTTGCCGATTTCGGAAC | Genotyping |
| CITRINE-mPLSP1 | Forward | GAGCTATACAAGGAGACAACGAAG | Genotyping |
| NOS-terminator | Reverse | AGACCGGCAACAGGATTCAATC | Genotyping |
| PLSP1-C166A | Sense | GGTGAGTTATTATTTCAGGAAGCCTgcaGCAAATGATATTGTC<br>PstI site underlined | Site-directed mutagenesis |
| PLSP1-C166A | Antisense | GACAATATCATTTGCtgcAGGCTTCCTGAAATAATAACTCACC<br>PstI site underlined | Site-directed mutagenesis |
| PLSP1-C286A | Sense | GCGGGACAGTGCTAGAAGGTGGCgcaGCTGTGGATAAGCAA<br>PvuII site underlined | Site-directed mutagenesis |
| PLSP1-C286A | Antisense | TTGCTTATCCACAGCtgcGCCACCTTCTAGCACTGTCCCGC<br>PvuII site underlined | Site-directed mutagenesis |
| PLSP1-S142A | Sense | CCAAGATATATTCCTTCTTTGgCTATGTATCCTAC | Site-directed mutagenesis |
| PLSP1-S142A | Antisense | GTAGGATACATAGcCAAAGAAGGAATATATCTTGG | Site-directed mutagenesis |
| T7-mPlsp1 | Forward | ATGGCTAGCATGACTGGTGGACAGCAAATGGGTGAGACAACGAAGTCT | Cloning |
| tpPlsp1-T7 | Reverse | ACCCATTTGCTGTCCACCAGTCATGCTAGCCATACTTGAATCCTTAAT | Cloning |
| attB1 | Forward | GGGGACAAGTTTGTACAAAAAAGCAGGCTCGCCCATG | GW Cloning |
| attB2 | Reverse | GGGGACCACTTTGTACAAGAAAGCTGGGTA | GW Cloning |

### References
